# Single-cell profiles of retinal neurons differing in resilience to injury reveal neuroprotective genes

**DOI:** 10.1101/711762

**Authors:** Nicholas M. Tran, Karthik Shekhar, Irene E. Whitney, Anne Jacobi, Inbal Benhar, Guosong Hong, Wenjun Yan, Xian Adiconis, McKinzie E. Arnold, Jung Min Lee, Joshua Z. Levin, Dingchang Lin, Chen Wang, Charles M. Lieber, Aviv Regev, Zhigang He, Joshua R. Sanes

## Abstract

Neuronal types in the central nervous system differ dramatically in their resilience to injury or insults. Here we studied the selective resilience of mouse retinal ganglion cells (RGCs) following optic nerve crush (ONC), which severs their axons and leads to death of ~80% of RGCs within 2 weeks. To identify expression programs associated with differential resilience, we first used single-cell RNA-seq (scRNA-seq) to generate a comprehensive molecular atlas of 46 RGC types in adult retina. We then tracked their survival after ONC, characterized transcriptomic, physiological, and morphological changes that preceded degeneration, and identified genes selectively expressed by each type. Finally, using loss- and gain-of-function assays *in vivo*, we showed that manipulating some of these genes improved neuronal survival and axon regeneration following ONC. This study provides a systematic framework for parsing type-specific responses to injury, and demonstrates that differential gene expression can be used to reveal molecular targets for intervention.

## Introduction

Insults to the central nervous system (CNS), whether acute (e.g. traumatic injury) or chronic (e.g. neurodegenerative disease), typically lead to irreversible damage. Some neurons die, and those that survive generally fail to grow new axons and re-establish synaptic connections. A well-recognized but poorly understood characteristic of these phenomena is that specific neuronal types are disproportionately affected even though causative insults are widely shared. For example, both huntingtin *(HTT)* and alpha-synuclein *(SNCA)* are broadly expressed in neurons, but mutations in *HTT* lead to Huntington’s disease with striatal GABAergic neurons as a main target, while mutations in *SNCA* lead to Parkinson’s disease with basal ganglia dopaminergic neurons as a main target (Fu et al., 2018; Saxena and Caroni, 2011). Similar differential effects have been documented for Alzheimer’s disease, Amyotrophic Lateral Sclerosis, and traumatic injuries to peripheral nerves and spinal cord (Conta Steencken et al., 2011; Welin et al., 2008). Thus, selective neuronal vulnerability is a common feature of neuronal insult.

We reasoned that comparing patterns of gene expression among neuronal types that are similar in many respects but differ in vulnerability might provide a means of pinpointing molecular pathways that contribute to their resilience. Although seldom used (Duan et al., 2015; Kaplan et al., 2014), this approach could complement previously employed target identification strategies that involve comparing neurons from different ages (e.g., regenerative developing vs. nonregenerative adult neurons; (Maclaren and Taylor, 1997)), regions (e.g., regenerative peripheral vs. nonregenerative central neurons;(Huebner and Strittmatter, 2009), or species (e.g., regenerative fish vs. nonregenerative mouse neurons; (Kizil et al., 2012)).

To explore this strategy, we analyzed the responses of mouse retinal ganglion cells (RGCs) to optic nerve crush (ONC), a long-studied model of traumatic axonal injury (Aguayo et al., 1991). RGCs are glutamatergic neurons that receive visual input from retinal interneurons and send their axons through the optic nerve, conveying visual information to retinorecipient areas in the brain (**Figure 1A**; Sanes and Masland, 2015). ONC transects RGC axons, causing the death of ~80% of RGCs within 2 weeks and ~90% within a month. Few survivors are capable of regenerating axons, but some can be provoked to do so by a variety of interventions, although none to date have proven capable of restoring useful vision (Benowitz et al., 2017).

**Figure 1.**
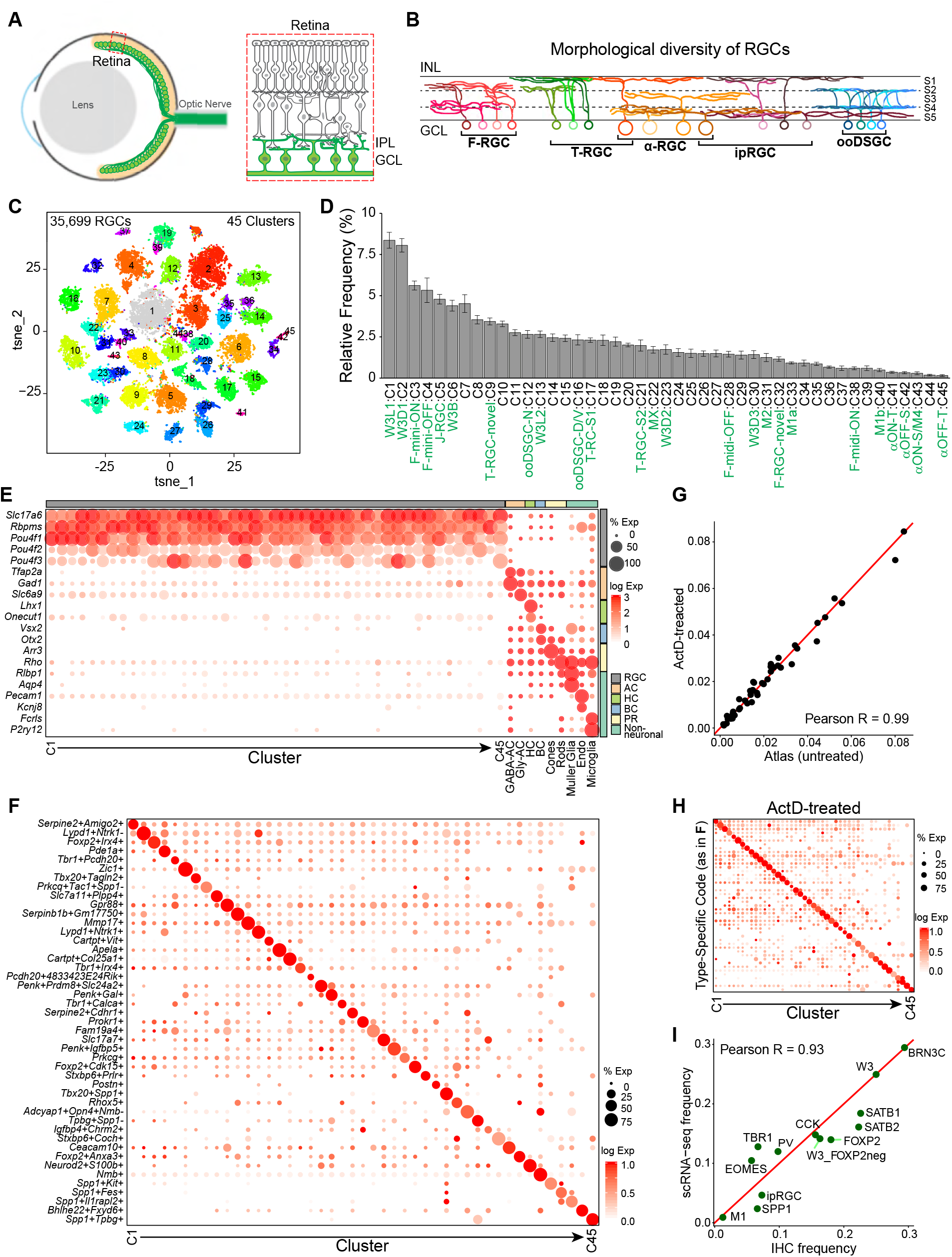
scRNA-seq reveals 45 molecularly distinct RGC types in adult mice. A. RGCs (green) reside within the innermost layer of the retina, the ganglion cell layer (GCL). Their axons bundle together to form the optic nerve. IPL, inner plexiform layer, GCL, ganglion cell layer. B. Dendrites of different RGC types have distinct lamination patterns within sublaminae (S)1 – 5 of the IPL, which determines their choice of presynaptic partners. Stereotyped morphologies are illustrated here for several RGC subclasses and types. INL, inner nuclear layer. C. t-distributed stochastic neighbor embedding (t-SNE) visualization of the transcriptional heterogeneity of 35,699 adult mouse RGCs. Cells are colored by cluster assignments, determined using graph clustering. Clusters are numbered in order of decreasing frequency. D. Relative frequencies of RGC clusters C1-45 (mean ±SD, n=10 replicates). Clusters that matched to known types or subclasses are labeled. E. Dotplot showing the expression patterns of marker genes (rows) specific to different retinal classes across RGC and non-RGC clusters in the data (columns; see color bars, top and right). The size of each circle is proportional to the percentage of cells expressing the gene, and the color depicts the average normalized transcript count in expressing cells. GABA-AC and Gly-AC, GABAergic and glycinergic amacrine cells; HC, horizontal cells; BC, bipolar cells; PR, photoreceptors; Endo, endothelial cells. F. Dotplot showing gene combinations (rows) that uniquely mark RGC clusters (columns). Representation as in panel E for single genes, here normalized to 1. 2- or 3-marker codes, always involve the presence of a marker A, and the presence (e.g. *A+B+* or *A+B+C+)* or absence (e.g. *A+B-*, or *A+B-C+)* of markers *B* and *C*. In such cases, the size of the circle indicates the percentage of cells satisfying the expression pattern, and the color depicts the average transcript count of positive markers in the cells, normalized to 1 for each combination. G. RGC type frequencies are highly similar between ActinomycinD (ActD)-treated (y-axis) and atlas (x-axis) retinas. H. Dotplot showing gene combinations that uniquely define each RGC type in nominal controls (as in **F**), are preserved in ActD-treated retinas. Row and column order as in **F**. I. Scatter plot showing tight correspondence (R_*Pearson*_ = 0.93) between relative frequencies of RGC groups found by scRNA-seq (y-axis) versus IHC (x-axis).

Several features make ONC an ideal model to study differential vulnerability across neuronal types: (1) All and only RGC axons pass through the optic nerve, so the timing and location of injury is precisely controlled, simultaneous and specific. (2) RGC loss occurs evenly across the retina so RGCs that live and die share the same microenvironment. (3) Although all RGCs share numerous features, they comprise >40 discrete types in mice, each with distinct morphology, connections and responses to visual stimuli (**Figure 1B**, and see below). (4) Some RGC types were recently shown to differ in their ability to survive or regenerate axons following ONC (Duan et al., 2015; Norsworthy et al., 2017; Perez de Sevilla Muller et al., 2014).

To survey the resilience of RGC types, we used single-cell RNA-seq (scRNA-seq), which we have previously applied to classify neurons in mouse and macaque retina (Macosko et al., 2015; Peng et al., 2019; Shekhar et al., 2016). We first generated a comprehensive atlas comprising 46 molecularly distinct types from >35,000 normal adult RGC transcriptomes, and used histological approaches to relate transcriptomic clusters to known and novel RGC types. Using this atlas as a foundation, we combined scRNA-seq with a novel computational approach to survey resilience among ~65,000 RGCs at 6 time points between 12 hours and 2 weeks after ONC. We determined the survival and kinetics of loss for each RGC type, finding dramatic differences, and assessed physiological and morphological changes that precede death. We then analyzed expression differences among RGC types before and following ONC, identifying genes that correlated with resilience or susceptibility. Finally, we used loss- and gain-of-function methods *in vivo* to test a subset of these genes, identifying some that regulate RGC survival and/or axon regeneration. Taken together, our work establishes a comprehensive molecular atlas of adult mouse RGCs, documents cellular and molecular changes preceding degeneration, and demonstrates that this approach can be leveraged to identify novel neuroprotective mechanisms.

## Results

### An atlas of molecularly defined RGC types

To generate a molecular atlas of RGC types, we isolated RGCs from adult (postnatal day [P]56) mice by fluorescence activated cell sorting (FACS), and profiled them by droplet-based scRNA-seq. Computational analysis of 35,699 high-quality single cell transcriptomes revealed 45 molecularly distinct clusters (**Figure 1C**), ranging in frequency from 0.15% to 8.4% (**Figure 1D**). All clusters expressed pan-RGC markers such as *Slc17a6* (which encodes the transporter VGLUT2), *Rbpms*, and at least one of the three *Pou4 (Brn3)* transcription factors, whereas markers of other retinal classes were present at low levels (**Figure 1E**). A few clusters could be matched 1:1 to previously characterized types based on differential expression (DE) of a single gene (e.g., *Jam2* for J-RGCS [Kim et al., 2008] and *Mmp17* for N-ooDSGCs [Kay et al., 2011]), but for most clusters, unique identity was conferred only by two-marker combinations (**Figure 1F**).

Since ~3 hours elapse between enucleation and RNA capture, we considered that clustering could be influenced by post-mortem alterations to gene expression, rather than intrinsic signatures. To test this possibility, we analyzed ~11,800 RGCs from retinas treated with actinomycinD (ActD) immediately upon enucleation, in order to block new transcription (Hrvatin et al., 2018; Wu et al., 2017). Although some differences were observed, such as the expected upregulation of immediate early genes (IEGs) in untreated retinas, the frequency of types, and their distinguishing markers were identical between Act-treated and untreated RGCs (**Figures 1G,H and S1A,B**).

Clusters were highly reproducible across biological replicates and three different computational approaches (**Figure S1C-E**). The number of estimated molecular types agree well with recent studies based on physiological, morphological and molecular methods (Baden et al., 2016; Bae et al., 2018; Rheaume et al., 2018) (**Table S1 and Figure S1F**). Finally, we compared the relative frequencies of RGC groups labeled immunohistochemically (IHC) in retinal whole mounts to their frequencies in the scRNA-seq data and found a striking correspondence (**Figure 1I**). Together, these results indicate that our atlas is comprehensive.

### scRNA-seq clusters correspond to morphologically-defined RGC types

To assess the morphology of molecularly-defined RGCs, we applied *in situ* hybridization and IHC to transgenic retinas in which RGCs were sparsely labeled (YFP-H line; ~200 RGCs per retina; (Samuel et al., 2011). We chose genes expressed by one or a few clusters, allowing us to validate novel markers for known types and characterize potentially novel types. For example, novel RGC clusters C10 and C24, which specifically expressed *Gpr88* and *Fam19a4*, respectively, possessed dendrites that were bistratified in sublaminae (S)2 and S4 of the inner plexiform layer (IPL), while dendrites of C25, which was labeled by the vesicular glutamate transporter-1 (VGLUT1) encoding gene *Slc17a7*, stratified exclusively in S5 (**Figure 2A**). (We use the convention of dividing the IPL into 5 sublaminae, S1-5; see **Figure 1B**). Other examples are shown in **Figure S2A-E** and results are summarized in **Table S2**.

**Figure 2.**
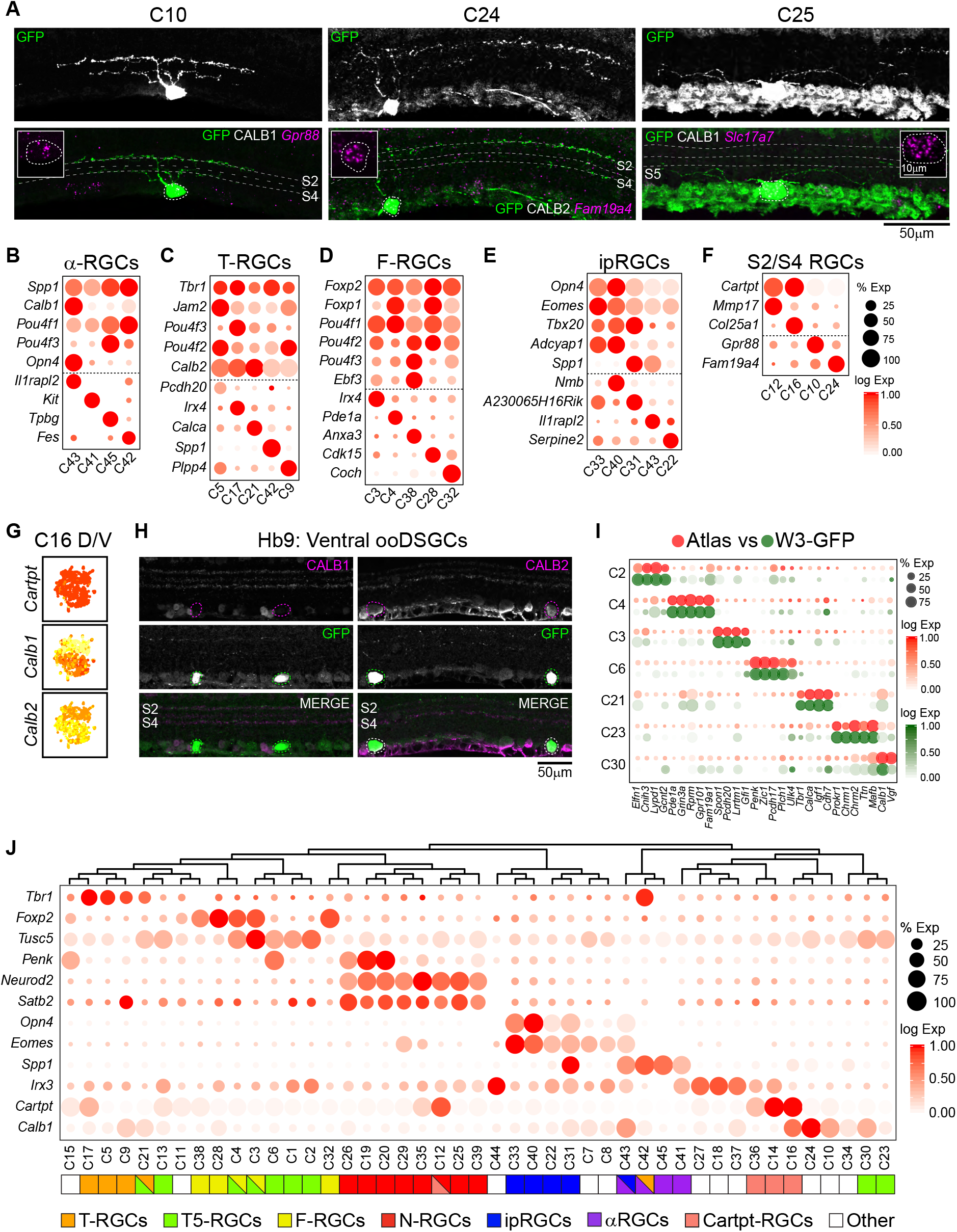
Correspondence of scRNA-seq clusters to RGC types. A. Characterization of novel RGC types by combining FISH (magenta) and IHC on sparsely labeled RGCs in the YFP-H line (green). Examples of S2/S4 laminating C10 and C24 RGCs expressing *Gpr88* (left) and *Fam19a4* (middle), respectively, and an S5 laminating C25 RGC expressing *Slc17a7* (right). IPL sublaminae are drawn based on CALB1 or CALB2 staining (white dashed lines). B-F Dotplots highlighting transcriptional distinctions among RGC types within subclasses. Dotted lines separate previously described markers (above) from novel markers identified in this study (below). B: αRGC types. C: T-RGC types. D: F-RGC types. E: ipRGC types. F: S2/S4 laminating RGC types. G. C16 comprising D/V-ooDSGCs can be partitioned into *Calb1+* (putative D-ooDSGCs) and *Calb1-* (putative V-ooDSGCs) cells. H. Consistent with the interpretation in panel G, GFP+ cells in the *Hb9* mouse line, which labels V-ooDSGCs, are CALB1- and CALB2+ (magenta). I. Dotplot showing consistent patterns of DE gene expression between W3 types (rows) detected in the droplet-based scRNA-seq atlas (red) and plate-based data from FACS-sorted W3 RGCs (green). Labeled by atlas cluster id. J. Transcriptional relatedness of RGC clusters visualized as a dendrogram reveals subclasses of RGC types (annotation bar, bottom). Dotplot shows expression of key subclass-enriched or -defining genes (rows) in clusters (columns).

### Transcriptome-assisted division of RGCs into subclasses

We and others have previously defined several groups of related RGC types, which we call subclasses. They include αRGCs, which express *Spp1* (osteopontin); T-and F-RGCs, defined by expression of the transcription factors *Tbr1* and *Foxp2*, respectively; ooDSGCs, defined by their physiological properties and bistratified dendrites; and intrinsically photosensitive RGCs (ipRGCs), defined by expression of *Opn4* (melanopsin) (Krieger et al., 2017; Liu et al., 2018; Rousso et al., 2016; Schmidt et al., 2011; Vaney et al., 2012) (Table S2). Subclasses exhibit some molecular overlap (for example ON- and OFF-sustained αRGCs express *Opn4* and *Tbr1*, respectively) but are largely distinct.

By double-labeling retinas for a defining subclass marker and a novel cluster-specific marker, we validated gene combinations that distinguish RGC types within each of these subclasses. For αRGCs (C41-43,45), the novel marker combinations of the 4 types are substantially more selective than those found previously by a candidate approach (Krieger et al., 2017) (**Figures 2B, S2A**). Our previous studies identified 4 T-RGC and 4 F-RGC types, but this new approach revealed a fifth type within each subclass (C9 and C32, respectively; **Figure 2C,D, S2B**). For ipRGCs, we discovered markers for M1 (C40), M2 (C31) and M4 (C43) types, including some that divide M1-RGCs (identified by high levels of *Opn4* and expression of *Adcyap1;* (Hannibal et al., 2002)) into two types (M1a, M1b), as well as an additional cluster (C22) that could correspond to the morphologically and physiologically characterized M3, M5 or M6 types (**Figure 2E**) (Quattrochi et al., 2019; Schmidt et al., 2011). For ooDSGCs, most of which are *Cartpt+*, we identified the nasal-preferring type (N-ooDSGC) by expression of *Mmp17* but only a single cluster (C16) expressed *Col25a1*, a marker of both D- and V-ooDSGCs (Kay et al., 2011). However, supervised analysis split this cluster with *Calb1* and *Calb2* in largely nonoverlapping cells, bringing the total number of RGC types to 46. Labeling of a line that marks only V-ooDSGCs confirmed that CALB1-positive cells are D-ooDSGCs and CALB2-high cells are V-ooDSGCs (**Figure 2F-H**).

Another potential subclass is defined by the transgene TYW3, which exhibits insertion site-dependent expression in several types of RGCs, one labeled brightly (W3B) and the others dimly (W3D), all of which share dendritic lamination in the middle third of the IPL (**Figure S2F**) (Kim et al., 2010; Laboulaye et al., 2018) (Krishnaswamy et al., 2015; Zhang et al., 2012). We isolated W3-RGCs by FACS and profiled them using Smart-seq2, to obtain deeper sequencing coverage (Ding et al., 2019; Picelli et al., 2013). Of 341 RGCs, 97% matched to 1 of 6 types in the atlas: W3B (identified by high expression of Sdk2; (Krishnaswamy et al., 2015)), F-mini-ON, F-mini-OFF and three others that we call W3D1-3. The remaining 3% corresponded to T-RGC-S2 (**Figure S2G,H**). Interestingly, all of these types expressed the integral membrane protein *Tusc5/Trarg1* (**Figure 2J**). Two additional atlas clusters, C1 and C13, were transcriptionally proximate to these types and *Tusc5/Trarg1*-positive; we call them W3-like (W3L) 1 and 2 (**Figure 2J**). This congruence identifies 9 types as members of a subclass that we provisionally call T5-RGCs. It includes 5 of the 6 most abundant RGC types and altogether accounts for approximately 40% of all RGCs.

Collectively, the subclasses defined above account for 26/45 RGC types, with each type occupying no more than two subclasses. Types within a subclass were usually but not always closely related molecularly: 4/5 T-RGCs (Tbr1+), 4/5 F-RGCs *(Foxp2+)*, 5/9 T5-RGCs *(Tusc5+)*, 4/5 ipRGCs *(Opn4+)*, and 3/4 αRGCs *(Spp1+)* were close relatives based on a hierarchical clustering analysis (**Figure 2J**). Our dataset also enabled the identification of putative novel subclasses based on transcriptional similarity and molecular markers. For instance, 8 closely related types co-expressed the transcription factors *Neurod2* and *Satb2* (provisionally N-RGCs; **Figure 2J**). It is tempting to speculate that types within this group, 7/8 of which are apparently novel, could also share cellular characteristics. The remaining 11/45 types were not assigned to a subclass due to the lack of a marker shared with proximal clusters; but they do exhibit some intriguing transcriptome-wide relationships to other types (**Figure 2J**). For example C7 and C8 are proximate to the known ipRGCs and express *Opn4* at low levels; they could also be ipRGCs. C10 and C24 are transcriptionally proximate to D/V-ooDSGCs (C16) and, like known ooDSGCs, are S2/S4 laminating (**Figure 2A**); they are candidates for the temporal-preferring (T) ooDSGC type.

### RGC types vary dramatically in susceptibility to ONC

Using the adult RGC atlas as a foundation, we assessed the resilience of types to ONC (**Figure 3A**). To this end, we profiled ~8,500 RGCs 14 days post ONC (dpc), at which point ~80% had died. Extensive injury-related changes in gene expression initially limited our ability to classify surviving RGCs to types, with only ~34% of 14dpc cells confidently mapping to types in the atlas using a “one-step” supervised classification framework (**Figure 3C**). We therefore formulated an alternative approach, leveraging data from RGCs collected at 5 intermediate time points. In this approach, transcriptomic signatures of RGC types were redefined at each time in order to assign cells to types at the next time point (**Figure 3B**). This allowed us to disambiguate gradual injury-related “state” changes for each RGC from its intrinsic type-specific signature. RGCs were assigned to types using a hybrid algorithm that combines supervised classification using gradient-boosted trees (Chen and Guestrin, 2016) and graph-based voting; we call the overall approach iterative-GraphBoost (iGraphBoost; see Methods).

**Figure 3.**
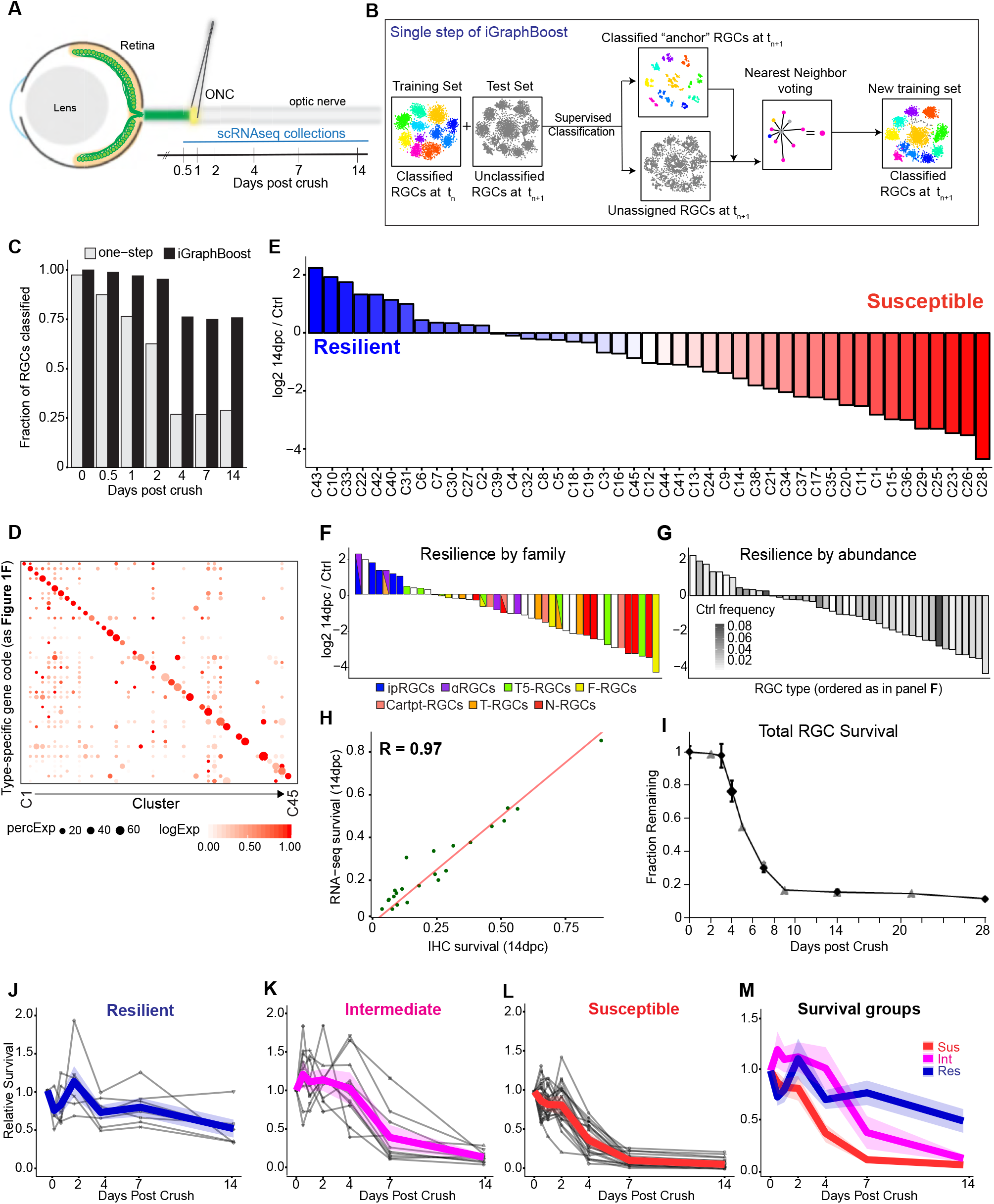
scRNA-seq profiling of RGCs following ONC. A. scRNA-seq was performed on RGCs collected before and at six times following ONC. 8,456-13,619 RGCs were collected at each time point. B. Illustration of a single step of the iGraphBoost procedure to classify RGCs collected at time t_n+1_ based on an atlas of RGC types at the previous time point t_n_. The procedure is initiated with Atlas RGCs at t_0_. In **Step 1**, gradient boosted trees trained on t_n_ RGC types are used to classify t_n+1_ RGCs. Only high-confidence assignments are applied, and a large number of RGCs remain unclassified at this stage. In **Step 2**, a Jaccard-weighted k-nearest neighbor graph built on all t_n+1_ RGCs is used to propagate labels via nearest-neighbor voting to unassigned RGCs, using the classified RGCs in step 1 as anchors. Successfully classified t_n+1_ RGCs are used to classify t_n+2_ RGCs in the next iteration. C. Fraction of RGCs that can be confidently assigned to types (y-axis) at each time point following ONC (x-axis). The “one-step” approach (grey) using the atlas RGCs as training data results in a significantly lower proportion of assigned cells among late injured RGCs compared to iGraphBoost (black). D. Dotplot showing that gene combinations uniquely defining each RGC type (row and column order as in **Figure 1F**) are maintained in 14dpc assigned by iGraphBoost, though reduction in expression level of some markers was observed. E. RGC type-specific resilience at 14dpc relative to control (Ctrl) rank ordered based on decreasing values of the relative frequency ratio at 14dpc vs. Ctrl. RGC types exhibit a wide spectrum of survival at 14dpc ranging from 1-98%. F. 14d survival ranking (as in E) colored by RGC subclasses. Overlapping subclasses are denoted by two-tone color bars. G. 14d survival ranking (as in E) colored by relative abundance in control. H. Scatter plot showing correspondence between the 14dpc survival rates of RGC groups as determined by scRNAseq and IHC (R_*Pearson*_ =0.97). 26 combinations of antibodies and transgenic lines (Table S3) were used label groups of RGC types covering a broad frequency range. I. Loss of RGC somas as determined by IHC for RBPMS in this study (diamonds; see **Figure S4F** for example images) or by retrograde labeling from superior colliculus (triangles; redrawn from (Galindo-Romero et al., 2011)). J.-M. Each RGC type can be assigned to one of three survival groups based on the pattern of cell loss across time. Individual graphs of relative survival, defined as the fraction of cells surviving at each time point, shown for 7 resilient types (J), 11 intermediate types (K) and 27 susceptible types (L), (see also **S3G**). Fluctuations in sampling frequency resulted in relative survival values >1 through 2dpc (where there is little death) for rare RGC types (frequency < 0.5%). Error bars are not included for individual types in panel **K-L** for clarity of presentation. Grey lines, relative survival for each type within the survival group; colored lines, mean relative survival across types; shaded ribbons, standard deviation of relative survival values across types. Fluctuations observed through 2dpc were within expected error (colored ribbons), in contrast to later time points. Solid lines, mean relative survival across types within a survival group; shaded ribbons, standard deviation. Group means are superimposed in **M**.

iGraphBoost assigned 89% of total injured RGCs to types, including 77% at 14dpc (**Figures 3C**). RGCs mapped by iGraphBoost maintained specificity of expression of the one and two-marker gene combinations through 14dpc (compare **Figures 1F and 3D**). To visualize the injured RGCs in a 2D representation we combined Liger, which utilizes non-negative matrix factorization (Welch et al., 2019), and t-SNE (**Figures S3A**). Encouragingly, clusters identified using the Liger representation were associated more strongly with type-specific identities assigned by iGraphBoost, than with other metadata such as time, mouse strain, or collection (**Figures S3C**). Nonetheless, some surviving RGCs could not be confidently classified by iGraphBoost. The proportion of “unassigned” cells increased over time, consistent with the idea that injury related transcriptional changes mask type-intrinsic signatures (**Figures S3D-F**).

Using results from iGraphBoost, we ranked RGC types by their frequency at 14dpc compared to control, a measure of resilience (**Figure 3E**). Type-specific survival rates varied continuously from ~1 % to ~98% at 14dpc. We refer to the 7 types that showed an increase in relative frequency >2-fold as ‘resilient’ (resRGCs); these types accounted for 8.1% of RGCs in control and 25.6% at 14dpc. Because some differences in apparent resilience could result from biases in collection, loss of particularly fragile cells, or misassignment by our computational approach, we also assessed survival for selected types using IHC (**Figures S4A-E, Table S3**). Histologically and transcriptionally derived frequencies were as highly correlated (Pearson r=.97) at 14dpc (**Figure 3H, Table S3**), as in controls (**Figure 1I**). Taken together, these data provide a comprehensive catalog of type-specific vulnerability of RGCs to injury.

We then asked whether relative resilience of RGC types correlated with overall molecular relationships. In some cases, correspondence was striking. For example, all ipRGC *(Opn4+)* types were resilient, and all N-RGC types were susceptible (**Figure 3F**). Other transcriptionally defined groupings of RGCs, however, contained types that differed greatly in resilience. For example, among αRGCs, which had previously been characterized as a resilient subclass (Duan et al., 2015), the two sustained types (C42, 43) were highly resilient but the two transient types (C41, 45) were relatively susceptible, despite clustering together transcriptionally (**Figures 3F, S4A-B**). Likewise, both rare and abundant types could be either resilient or vulnerable (**Figure 3G**). Thus, transcriptional proximity and frequency are imperfect predictors of resilience.

### Dynamics of RGC survival after injury define three survival groups

Few RGCs die during the first 3 days after ONC, ~70% die over the next 5 days, and numbers then decline gradually to ~10% survival at 28dpc (**Figures 3I, S4F**). Based on their kinetics of loss, RGC types could be partitioned into three survival groups: the 7 resilient types (8.1% of control RGCs) declined gradually, reaching ~50% survival at 14dpc; 11 “intermediate” types (27.2% of control RGCs) exhibited a striking decline between 4 and 7dpc; and 27 susceptible types (64.7% of control RGCs) were already severely reduced by 4dpc (**Figures 3J-M**). Thus, the survival of the intermediate and susceptible RGCs differed dramatically at 4dpc (susRGCs: 39% ± 21%; intRGCs: 95% ± 25%). Unsurprisingly, these groups correlated well with rankings by survival at 14dpc alone (**Figure S3G**). As above, we validated scRNAseq-derived survival kinetics of RGC subclasses with distinct survival rates using IHC (**Figures S4F-H**), demonstrating good correspondence throughout the time course.

### Physiological characteristics of resilient and susceptible RGCs

The resilience of ipRGCs and sustained αRGCs suggested that resilient RGCs may share common functional properties. However, visual responses are currently unknown for most molecularly defined types. Therefore, we monitored physiological characteristics of individual injured RGCs over time, using our recently developed method for *in vivo* recording (Hong et al., 2018). Briefly, a flexible electro-recording mesh carrying 32 electrodes is injected intravitreally, where it coats the inner retina without disturbing normal eye function; spike sorting protocols identify up to 4 cells per electrode, and provide wave-form signatures that allow longitudinal tracking of the same cell over multiple recording sessions (**Figures 4A,B**).

**Figure 4.**
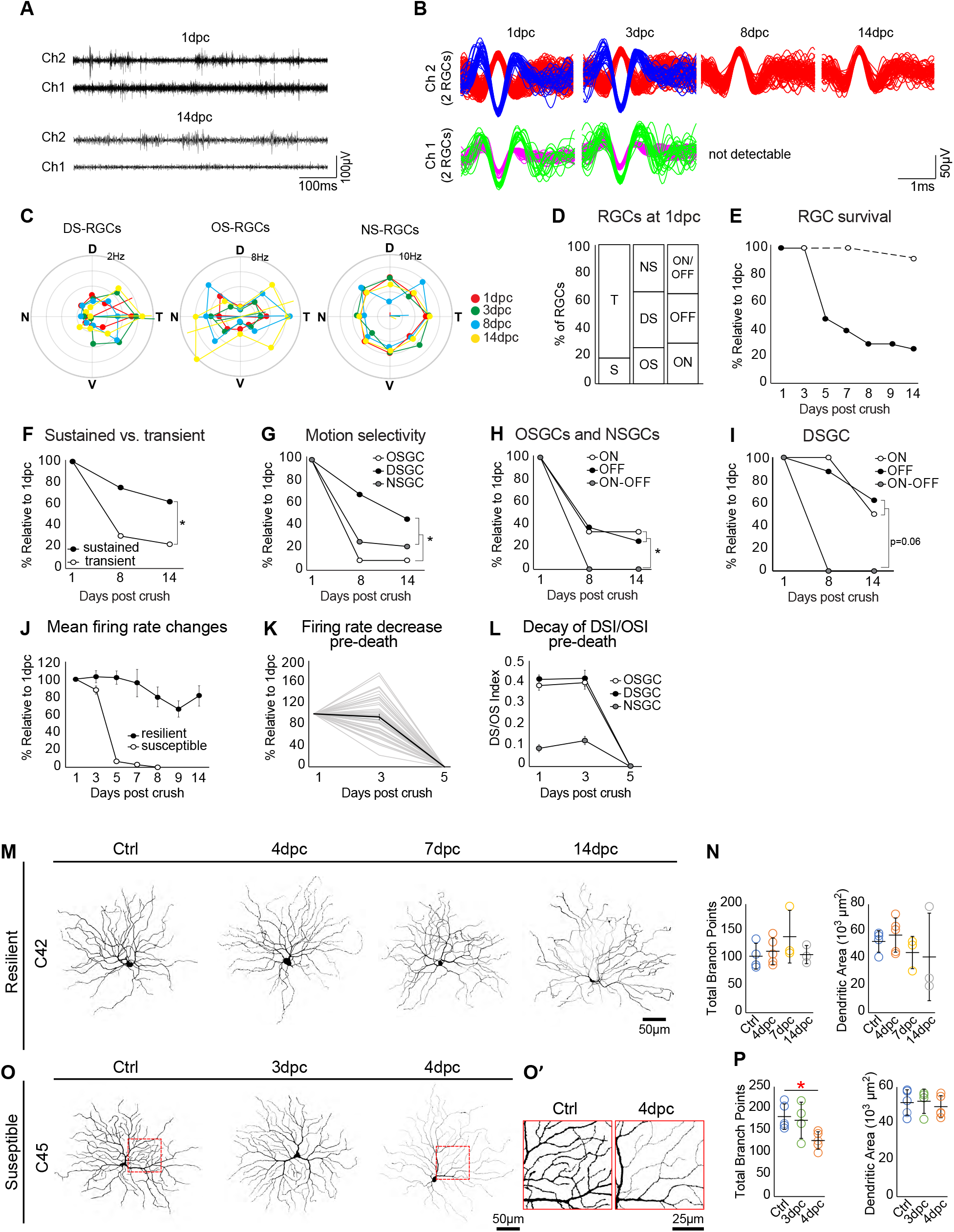
Physiological characteristics of resilient and susceptible RGCs. A. Representative recordings of two out of 32 channels in 1dpc and 14dpc mesh-implanted retinas. B. Sorted spike waveforms for two individual RGCs per channel (rows) represented in **A** recorded over multiple days. Ch1 shows spike waveforms of two sorted RGCs (purple and green lines) on 1dpc and 3dpc; cells have died by 8dpc. Ch2 shows waveforms of two sorted RGCs (blue and red) on 1dpc and 3dpc, but only one RGC was still detectable at 8dpc. C. Polar plots of responses of direction-selective (DS), orientation-selective (OS) and neither orientation-nor directions-selective (NS) RGCs to gratings moving in each of 8 directions. Each plot shows measurements from the same cell on different days. D. Proportion of RGCs by response type within each response category (columns) at 1dpc. S, sustained, T, transient, ON, OFF and ON/OFF, responds to light increments, decrements or both. E. RGC survival as a function of time in physiological recordings following ONC (black line) compared to uncrushed control (dotted line shows data replotted from (Hong et al., 2018)) F. Sustained RGCs survive better than transient RGCs as assessed by physiology (* = p<0.03 by Fisher’s Exact Test). G. OSGCs are more susceptible than DSGCs or NSGCs (* = p<0.04 by Fisher’s Exact Test). H. Among RGCs that are either OS or NS, ON-OFF cells are more susceptible than ON or OFF cells (* = p<0.03 by Fisher’s Exact Test). I. Among DSGCs, ON-OFF cells (ooDSGCs) are susceptible than ON or OFF cells (p=0.06 at 14dpc by Fisher’s Exact Test). J. Average firing rates for RGCs that survive until 14dpc or die by 8dpc. K. RGCs that are dead by 5dpc exhibit little changes in firing rate between 1-3dpc. L. RGCs that are dead by 5dpc exhibit little change in direction/orientation selectivity index (DSI/OSI) between 1-3dpc. M. *En face* morphology of resilient RGCs (αOFF-S, C42) at Ctrl, 4, 7, and 14dpc. N. Quantification of C42 morphological complexity (total branch points) and size (dendritic area) shows no significant difference between time points for either measure (one-way ANOVA with post-hoc Tukey HSD test). Data are shown as mean±SD. O. *En face* morphology of susceptible RGCs (αOFF-T, C45) at Ctrl, 3, and 4dpc. O’ showing zoomed in views of dendrites at Ctrl and 4dpc. P. Quantification of C45 morphological complexity as in N. * = p<0.04; one-way ANOVA with post-hoc Tukey HSD test. Data are shown as mean ±SD.

To measure light response properties, we implanted the mesh directly after ONC and recorded RGC activity every 1-2 days over periods of 6-14 days, obtaining longitudinal data from a total of 142 cells in 4 mice. We used full field illumination and gratings moving in each of 8 directions to determine whether RGCs exhibited responses that were sustained or transient; orientation-, direction-, or non-selective (OSGCs, DSGCs, and NSGCs, respectively; **Figure 4C**); and ON, OFF, or ON/OFF (firing in response to luminance increase, decrease or both). At 1dpc, we detected all these functional types, indicating that the method sampled RGC types broadly (**Figure 4D**).

We then tracked the survival of RGCs over 14 days to identify resilient and susceptible populations. We defined a cell as “dead” if its mean firing rate decreased below 0.5 Hz for at least two consecutive days. ~60% of RGCs died between 3 and 7dpc, with ~74% dead by 14dpc (**Figure 4E**). In contrast, less than 10% of cells were lost over two weeks of recordings from uninjured retinas (Hong et al., 2018), indicating that the loss reflects ONC-related death rather than recording instability. Moreover, the survival dynamics mirror those determined histologically (**Figure 3I**), suggesting that neurons are not silent for substantial periods prior to their death.

We used this method to ask whether surviving RGCs were enriched for specific response types. RGCs with sustained responses survived ~3-fold better than those with transient responses (**Figure 4F**), consistent with scRNA-seq results for sustained and transient αRGCs. Because αRGCs comprise <5% of all RGCs, the physiological result suggests that the relationship between sustained responses and resilience is a general one. OSGCs were more susceptible than DSGCs or NSGCs (**Figure 4G**). Resilience did not differ between ON or OFF types but ON-OFF types were more vulnerable, independent of feature selectivity (**Figure 4H**). This vulnerability is consistent with the known susceptibility of ooDGCS, which have transient responses, but did not extend to other DSGCs (**Figure 4I**). These results reveal a correlation between physiological properties and resilience.

Our longitudinal measurements also enabled the assessment of physiological changes preceding cell death. Overall firing rate of resilient RGCs (i.e., those detectable at 14dpc) varied little during the measurement period (**Figure 4J**). Similarly, for RGCs that died between 3 and 5dpc, the firing rate and orientation- and direction-selectivity indices were largely unchanged between days 1 and 3 (**Figures 4K,L**). These results suggest that RGCs maintain activity levels and presynaptic inputs, which determine response properties, until shortly before they die.

### Morphological changes in resilient and susceptible RGCs

The observation that functional responses of RGCs were retained until at least 48hrs prior to death raised the question as to whether their structural integrity was similarly maintained. We therefore tracked changes in dendritic morphology of 3 resRGC types (ipRGC M2, αRGC OFFS, and αRGC ON-S/ipRGCC M4) and 3 susRGC types (αRGC OFF-T, and 2 ooDSGC types) after ONC. We used sparse morphological labeling with IHC and dendritic lamination to identify types (**Figures S5A-C**), imaged them in whole mounts, and reconstructed their dendrites.

Strikingly, resRGCs maintained robust dendritic morphology through 14dpc, with no significant decrease in dendritic area or arbor complexity (**Figures 4M,N; S5D,E**). Along with functional results, this implies that resRGCs maintain their integrity after ONC. In contrast, susRGC types were scarce by 7dpc, consistent with their survival dynamics. Interestingly, however, all three susRGC types also maintained their dendritic area through 4dpc, though their dendrites often appeared thinner and fainter than those in controls and two of the three susRGC types exhibited a significant reduction in total branch points, a measure of dendritic complexity (**Figures 4O,P; S5F,G**). Together with physiological measurements, this morphological analysis raises the possibility that a substantial window exists during which surviving RGCs could be receptive to regenerative therapies.

### Global gene expression changes after ONC

To ask when injury response programs are activated in RGCs, we first characterized the dynamics of globally regulated gene expression after ONC, identifying 771 temporally DE genes that were broadly shared across types (**Table S4**). Genes were partitioned into 8 modules (Mod1-8) by k-means clustering (**Figure 5A**), identifying gene sets with distinct temporal dynamics that were enriched for different gene ontology (GO) biological processes (**Table S5**). For example, module 1 (Mod1), which comprised genes whose expression began to decline as early as 0.5dpc, was enriched in GO terms associated with functions carried out in healthy neurons such as action potential, synaptic vesicle exo/endocytosis, retrograde axon transport and microtubule polymerization (**Figure 5B**). In contrast, Mod5 and Mod6, comprising genes upregulated around 2dpc, were associated with apoptosis and stress pathways such as metabolic and ER stress, unfolded protein response, and catabolism (**Figure 5C**). Globally regulated genes generally did not show strong type-specific differences, with the notable exception of ipRGC types, which exhibited considerably lower upregulation of Mod5, 6 and 7 genes than other types (**Figures 5D-E, S6A-B**).

**Figure 5.**
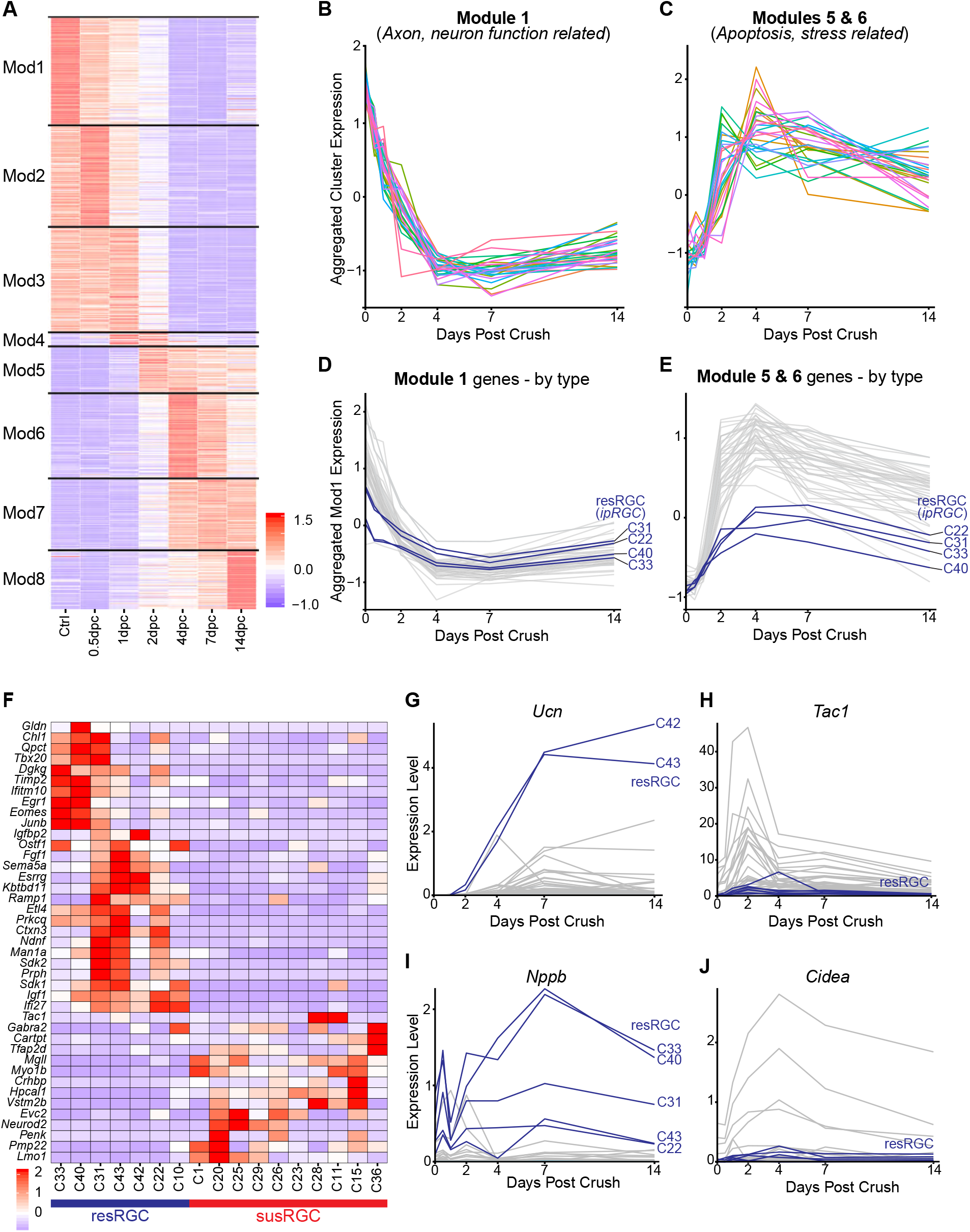
Global changes in gene expression following injury. A. Heatmap of genes showing temporal variation following ONC. Expression values of each gene (row) is averaged across all RGCs at a given time point (columns), and then z-scored across times prior to plotting. Black bars separate genes into 8 modules (Mod) based on temporal dynamics. B. Mean temporal dynamics of individual genes (lines) from Module 1 that were associated with GO biological processes related to axon and neuronal functions. Genes are listed in **Table S4**. C. As in **B**, for Modules 5 and 6 for genes associated GO biological processes related to apoptosis or various stress pathways. D. Expression dynamics of genes from B plotted for each RGC type (lines). Blue lines correspond to ipRGC types (C31, 22, 40, 33). Expression values for each type were z-scored to track relative changes. E. Same as **D**, for genes from **C** F. Expression patterns of DE genes (rows) distinguishing the 7 resRGC types and the 10 most susceptible RGC types (columns), based on 14dpc survival in the uninjured retina (**Figure 3F**). Values were z-scored along each row prior to plotting. G.-J. Averaged temporal dynamics of candidate genes selectively upregulated in resRGC or susRGC types (lines). Blue lines correspond the 7 resRGC types, including types that upregulate *Ucn* (C42, 43) or *Nppb* (C22, 31, 33, 40, 43) (left panels), which were not enriched for *Tac1* or *Cidea* (right panels).

### Gene expression correlating with resilience and vulnerability

Next, we next sought type-specific gene expression patterns that correlated with resilience or vulnerability. First, we compared baseline (control) transcriptomic profiles across the three survival groups (**Figure 3N**). Several genes were expressed in multiple resRGC types but showed little to no expression in susRGC or intRGC types (**Figures 5F, S6C**). Among them were two IEGs *(Junb* and *Egr1)*, enriched in ipRGCs. In light of the upregulation of IEGs by dissociation (**Figure S1F**), we verified that IEG and other type-specific expression patterns were maintained in RGCs treated with ActD, and are therefore likely to be intrinsic properties (compare **Figures 5F and S6D**). Notably, with the exception of *Igf1*, we found few DE genes that were enriched in all resRGCs but no susRGCs or vice versa, suggesting heterogeneity in factors in mediating resilience or susceptibility across types.

We also asked if resilient and susceptible RGC types up- or down-regulated different sets of genes following injury. As was the case for the analysis at baseline, few genes were up- or down-regulated in all resRGC or susRGC types, but many were upregulated selectively in several resRGC but no susRGC types or vice versa (**Figure 5G-J**). resRGC-enriched genes were generally maintained through 14dpc, suggesting they could play a role in long term survival. In contrast, expression of susRGC-enriched genes generally peaked at 2-4dpc then declined, coincident with the onset of degeneration, suggesting that their expression could be predictive of cell death.

### scRNA-seq-derived candidates promote neuroprotection of RGCs

Genes enriched in resRGCs included three previously described mediators of RGC survival and/or axon regeneration: *Igf1* (7/7 resRGCs), *Opn4* (5/7) and *Spp1* (3/7) (Duan et al., 2015; Dupraz et al., 2013; Li et al., 2016; Zhang et al., 2019). To ask whether genes selectively expressed in resilient or susceptible RGC types included additional factors that affected survival, we chose 10 candidates to test (**Table S6**). Adeno-associated viral serotype 2 (AAV2) vectors were used to overexpress (OE) genes correlating with resilience or to mutate (knock-out, KO) genes correlating with susceptibility. For KO experiments, we infected retinas from *LSL-Cas9* mice crossed to *Vglut2-Cre* mice (to express Cas9 in all RGCs) with AAV2 vectors encoding a single-guide RNA (sgRNA). We injected AAV intravitreally 14d prior to ONC to infect a high percentage of RGCs and quantified RGC survival by IHC at 14dpc (**Figure 7A, S7A**).

We began with a pair of genes that displayed intriguing expression patterns: urocortin *(Ucn)*, which encodes a peptide from the corticotropin-releasing factor family, and corticotropin releasing hormone binding protein *(Crhbp)*, a secreted glycoprotein that inhibits UCN-mediated activity (Seasholtz et al., 2002). *Ucn* was upregulated post-ONC in the two sustained αRGC types but not in other RGCs, while *Crhbp* was selectively expressed in multiple susRGC types (**Figure 6A,B**). The CRH receptor *(Crhr1)*, through which UCN signals, was broadly expressed among RGC types (**Fig. S7B**). We increased *Ucn* levels by AAV2-based OE or by injection of recombinant protein, and decreased *Crhbp* expression by AAV2-CRISPR-based KO with two different sgRNAs (**Figure S7A**) All four treatments significantly increased RGC survival (**Figure 6C,J, Table S6**).

**Figure 6.**
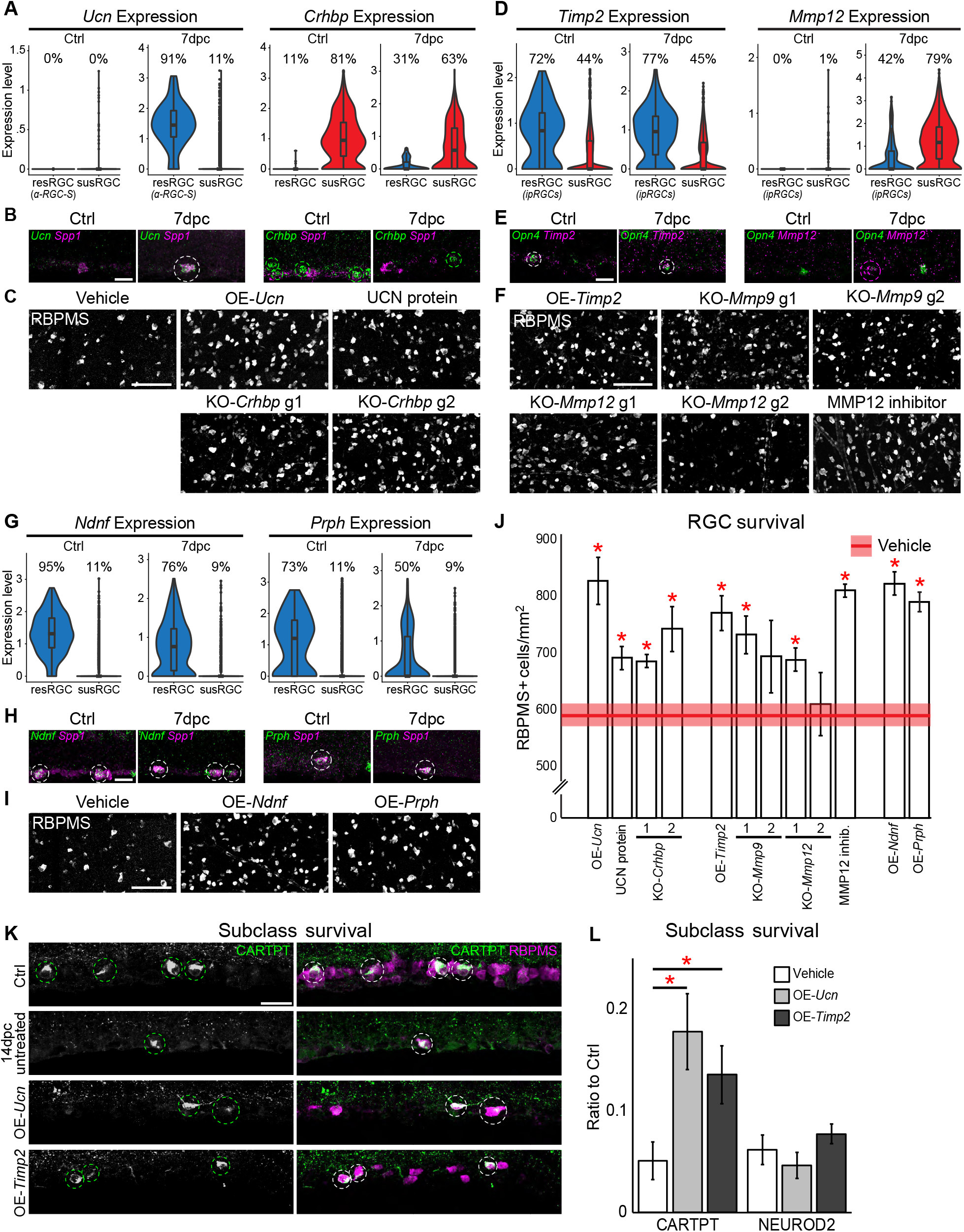
Genes that affect RGC survival. A. *Ucn* is selectively upregulated in sustained αRGC’s (α-RGC-S; C42, 43) and *Crhbp* is selectively expressed in a subset of susRGC types (C14, 15, 17, 24, 26, 28, 39). Violin plots show merged expression for indicated clusters at 0 and 7dpc. The number above the violins indicates the percentage of cells expressing the marker within each subset. Box plots depict the median and interquartile range. B. FISH of retinal sections shows *Ucn* upregulation at 7dpc in *Spp1+* RGCs (α-RGCs marker): white circles. *Crhbp* is expressed in a set of *Spp1-* RGCs (non-α-RGCs) before and after ONC: green circles. C. IHC in retinal whole mounts for RBPMS shows increased survival of RGCs at 14dpc following OE-Ucn, *KO-Crhbp*, or injection of UCN protein. D. *Timp2* is selectively expressed in the resilient ipRGCs (C22, 31, 33, 40, 43) before and after ONC. *Mmp12* is upregulated in a broad subset of susRGCs (C7, 8, 11, 12, 14, 17, 18, 23, 24, 27, 28, 43, 37, 39, 41) after crush but is low in ipRGCs in scRNAseq data. Violin plots as in **A**. E. FISH of retinal sections as in **B**. F. IHC in retinal whole mounts as in **C**. G. Expression in resRGC subsets at 0 and 7dpc of *Ndnf* (C22, 31, 43) and *Prph* (C31, 43). Violin plots as in panel **A**. H. FISH of retinal sections as in **B**. I. IHC of retinal whole mounts as in **C**. J. Total RGC survival (RBPMS+ cells; mean ± SEM) in whole mounts following interventions shown in **C, F**, and **I**. Red line and ribbon, mean RBPMS density ±SEM. *adjusted p-value <.05 (FDR). K. IHC showing increased survival of CARTPT+ RGCs (circles) at 14dpc following OE-Ucn and *OE-Timp2* compared to vehicle. Top row, CARTPT+ RGCs at 0dpc. L. IHC quantification showing selective survival of CARTPT+ RGCs (C12, 14, 16, 36) compared to NEUROD2+ RGCs (C12, 19, 20, 25, 26, 29, 35, 39) at 14dpc following indicated treatments. y-axis, #positive per section RGCs at 14dpc/control. Performed on retinal sagittal sections through the optic nerve. * p-value <.05 (FDR adjusted). Scale bar: 25μm for **B,D,H,K**; 100μm for **C,F,I**

A second pair of related genes was *Timp2*, an inhibitor of matrix metalloproteinases (MMPs) and *Mmp12*, a target of TIMP2 (Koppisetti et al., 2014). *Timp2* was selectively enriched in resilient ipRGCs at baseline and maintained its expression following ONC. Conversely, *Mmp12* was broadly upregulated after ONC but upregulation was particularly modest in ipRGCs (**Figure 6D,E**). AAV-OE of *Timp2* enhanced survival, as did AAV-KO of *Mmp12* with one of two sgRNAs (**Figure 6F, J, Table S6**). A small molecule inhibitor selective for MMP12 also improved survival. Because MMPs have overlapping functions, we surveyed the expression of *Mmp’s* in our scRNA-seq data and found that only *Mmp9* was expressed at a detectable level in multiple RGC types (**Figure S7C**). Targeting *Mmp9* with either of two sgRNAs also improved survival (**Figure 6F, J**), though the increase was only statistically significant for one sgRNA.

Of the other five genes tested, two selectively expressed by resRGCs improved survival: neuron-derived neurotrophic factor, *Ndnf*, which encodes a fibronectin III domain containing glycosylated secretory protein (Kuang et al., 2010), and peripherin, *Prph*, which encodes a type III neurofilament protein (Thompson and Ziff, 1989) (**Figure 6G-J, Table S6**). In contrast, KO of three genes enriched in subsets of susRGCs *(Evc2, Tac1* and *Hpcal1)* had no significant effect (**Figure S7D,E**).

Genes would be particularly useful targets if they were able to rescue neurons that do not express them endogenously. We asked whether the protective genes *Ucn* and *Timp2*, expressed by resRGCs, would improve survival of susceptible RGC types if expressed broadly. To test this idea, we used two markers (CARTPT, NEUROD2) that label susceptible RGC subclasses (**Figure 3H**), neither of which expresses *Ucn* or *Timp2* at high levels before or after ONC. Both OE-Ucn and *OE-Timp2* increased survival of CARTPT+ RGCs but not NEUROD2-RGCs (**Figure 6K,L**). Thus, these interventions can protect some but not all susceptible RGC types.

### Factors correlating with resilience also stimulate axon regeneration

While our screen was focused on neuroprotection, the targets we identified might also promote axon regeneration. To test this possibility, we anterogradely labeled RGC axons by intravitreal injection of fluorescently conjugated cholera toxin B subunit (CTB647) at 12dpc (**Figure 7A**). We collected optic nerves at 14dpc and quantified labeled axons at 500-2000μm from the crush site. OE-Ucn, UCN protein, *OE-Timp2, KO-Crhbp* and *KO-Mmp9* all promoted significant overall regeneration (**Figure 7B,C,E,F, Table S6**), with some regenerating axons extending >1500μm. In contrast, overexpression of *Ndnf* and *Prph*, showed minimal effects on regeneration (**Figure 7D,G**). These results encourage the hope that our screen will be useful for discovery of targets for axon regeneration as well as neuroprotection.

**Figure 7.**
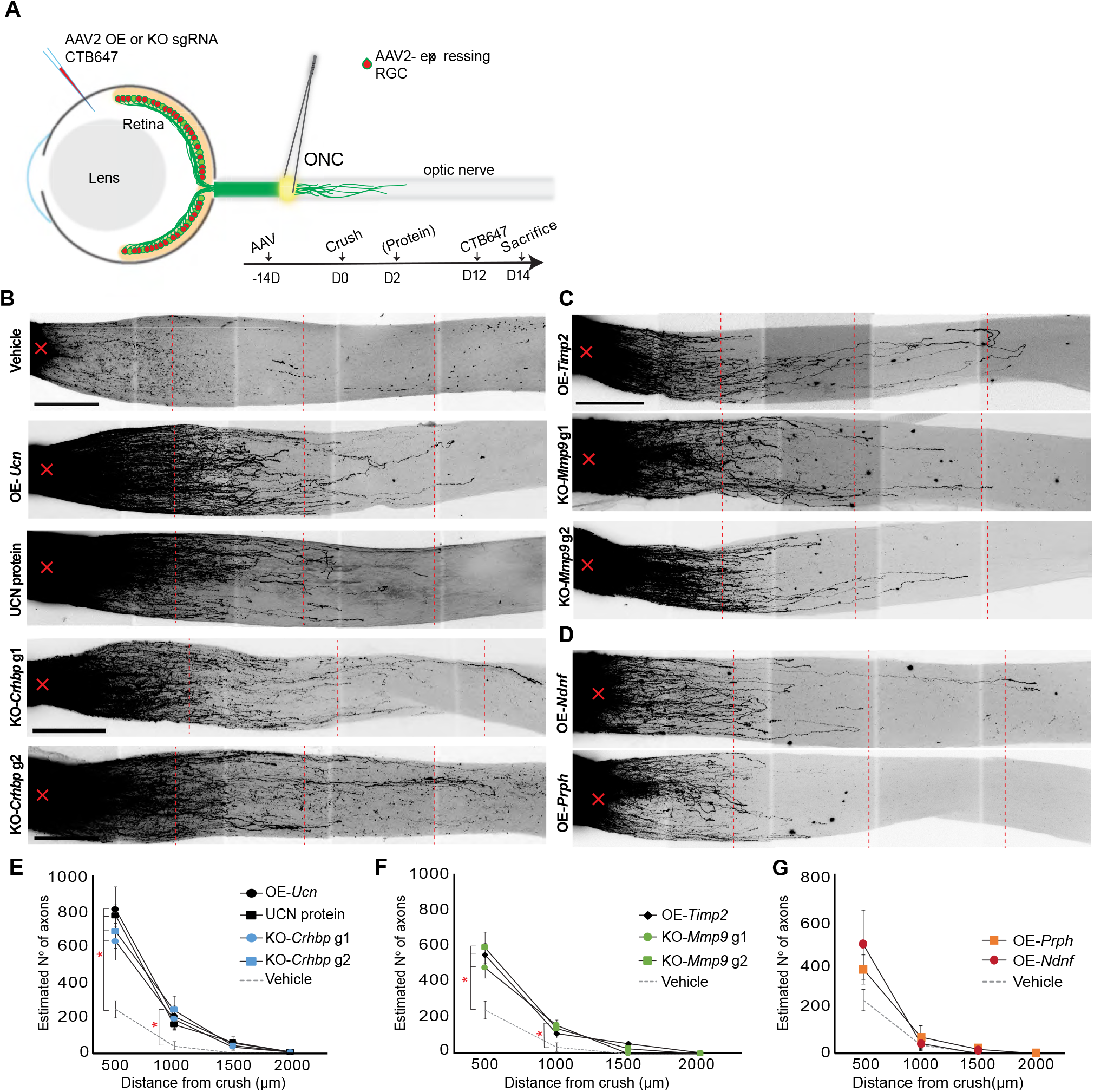
Genes that promote RGC axon regeneration. A. *In vivo* OE and KO. An AAV2 carrying the OE gene or KO sgRNA is injected intravitreally 14 days before the crush. At 12dpc regenerating axons are anterogradely labeled via CTB647 injection. UCN protein was injected at 2dpc. B. Maximum projections of cleared optic nerves showing anterograde-labeled RGC axons at 14dpc following vehicle injection or indicated treatment of Ucn (OE or protein) and KO-*Crhbp* (g1 and g2). C. Same as **B**, following *OE-Timp2* and *KO-Mmp9* (g1 and g2). D. Same as **B**, following *OE-Ndnf* and *OE-Prph*. E.-G. Quantification of axon regeneration. * p < 0.05 two-tailed Student’s t-test of area under the curve evaluated using numerical integration. 2-way ANOVA or mixed effects analysis with Bonferroni correction for individual distances are shown in **Table S6**. In **B-D**, Scale bar, 250μm; X = crush site; red lines – 500, 1000, 1500μm distances from crush site.

## Discussion

We generated a transcriptome-based atlas of adult mouse RGC types, and used it as a foundation to track type-specific responses to injury. We identified a spectrum of resilience among RGC types, and documented transcriptomic, physiological and morphological changes preceding degeneration. We then manipulated genes selectively expressed in resilient or vulnerable types, finding some that promote RGC survival and axon regeneration following ONC.

### An atlas of mouse RGC types

Analysis of 35,699 adult RGC transcriptomes revealed 45 cell clusters. Several lines of evidence indicate that the transcriptomically defined clusters correspond to cell types: (a) The number of types is similar to those from recent large-scale surveys based on serial section electron microscopy (≥35;(Bae et al., 2018), optical imaging of electrical activity (≥32; (Baden et al., 2016) and scRNAseq of neonatal retina (40; (Rheaume et al., 2018)). (b) Several clusters could be assigned to types based on previously known markers. (c) For others, *in situ* hybridization with genes identified from scRNA-seq allowed us to assign clusters to known types or find morphological signatures of previously uncharacterized molecular types. Thus, RGCs join retinal bipolar cells (Shekhar et al., 2016) as a second neuronal class for which transcriptomic criteria tightly correspond to types as defined by classical criteria. This encourages the belief that high-throughput molecular profiling methods, which are currently the most scalable, represent a reliable approach to categorize cell types in the mammalian nervous system.

We cannot, however, be sure that we have captured all RGC types for three reasons. First, our sample size permitted the identification of a type that comprises ~0.15% of all RGCs, but rarer types may have gone undetected. Second, cell dissociation can create biases and a fragile cell type could be missed. Third, V- and D-ooDSGCs formed a single cluster that we split by a semisupervised analysis, yielding a total of 46 RGC types from 45 cell clusters. These types are extremely similar; indeed, despite intensive study, no endogenous markers had been found that distinguish them (Kay et al., 2011). They do form separate clusters in early postnatal retina (I.E.W, K.S and J.R.S. in preparation), suggesting that distinguishing genes, presumably including those that specify their distinct connectivity, are likely downregulated in adulthood. Although supervised analysis of other clusters did not reveal additional subdivisions, we cannot rule out the possibility that some other closely related types also co-clustered.

The molecular atlas provided new insights into RGC subclasses. First, for several subclasses previously defined by morphological, physiological or genetic criteria, we identified novel members; for example, a fifth T-RGC and a fifth F-RGC. Second, members of a subclass generally showed a global transcriptomic relationship. For example, the T5-RGC types, 6 of which were previously shown to share dendritic lamination, were transcriptomically proximate (5/9 types) and shared expression of *Tusc5. Neurod2* and *Satb2* are co-expressed by 8 transcriptomically proximate types, 7 of which are novel. These and other novel types clustering near known types may share cellular features. On the other hand, types that share functional or structural characteristics were not always neighbors on the dendrogram (**Figure 2J**), particularly when a type was a member of multiple subclasses. For example, the *Neurod2+* ooDSGC C12 was transcriptomically distant from the *Neurod2^-^* ooDSGC C16.

### Resilience to injury varies among cell types

To characterize RGC survival after ONC, we applied three independent approaches – scRNA-Seq, IHC, and *in vivo* physiology. We found strong agreement among the criteria in their assessment of type-specific RGC resilience and draw four main conclusions. First, survival differed dramatically among types, from 1% to 98% over two weeks. Second, types that differed in overall resilience also differed in the time course of death, as epitomized by the three survival groups. Third, resilience was not “binary” but rather varied continuously across types.

Fourth, in some but not all cases, resilience correlated with molecular or physiological properties of RGC types. For example all 5 ipRGC types were resilient, extending previous reports on M1 and M4 αRGCs (Duan et al., 2015). Similarly, RGCs with sustained light responses outperformed those with transient responses. Conversely, all 8 N-RGC types survived poorly. Nonetheless, we did not find a single characteristic that predicted resilience. This could partially reflect technical limitations – for example, failure to detect lowly expressed genes, inadequacies of our analytical methods or post-transcriptional molecular determinant (e.g., post-translational modifications). However, we favor the idea that different resilient neuronal types utilize different survival strategies.

The resilience of sustained RGCs provides an intriguing parallel to patterns of motoneuron survival in Amyotrophic Lateral Sclerosis: motoneurons that innervate slow twitch muscle fibers (slow motoneurons) exhibit tonic (~sustained) activity and survive better than fast motoneurons, which exhibit phasic (~transient) activity (Pun et al., 2006). This correspondence suggests the possibility of a general association between firing pattern and resilience, the basis of which remains to be explored.

### Cellular changes in neurons with differential resilience

Knowing the kinetics of loss for each RGC type allowed us to characterize molecular, morphological and physiological alterations in the days prior to death. Our main result is that morphological and physiological changes were surprisingly mild up until shortly before somatic loss: dendritic morphology, firing rate and feature selectivity of RGCs remained at close to control levels until at least two days before death. This observation contrasts with models of glaucoma, where dendritic shrinking and functional decline prior to death is striking (Della Santina et al., 2013; Liu et al., 2011). They also seemingly contrast with other studies that observe significant dendritic shrinkage after ONC (e.g., (Agostinone et al., 2018)); however, those studies focused on broader subclasses of RGCs, so changes in dendritic area could reflect differential survival of RGC types within the subclass being assayed. It is also possible that our reliance on molecular marker expression excluded highly atrophied cells from our analysis.

### Genes that affect resilience and susceptibility

Although we did not find universal gene expression programs that predicted resilience, differences between resilient and vulnerable types, in controls or following ONC, led to identification of several promising candidate modulators of survival. Two intriguing pairs were identified in which a gene and its antagonist were expressed in opposing populations.

The first pair was *Ucn* and *Crhbp*, expressed by resRGC and susRGCs, respectively. UCN has been shown to promote survival of hippocampal and dopaminergic neurons, potentially by increasing intracellular cAMP levels (Abuirmeileh et al., 2007; Huang et al., 2011; Pedersen et al., 2002). CRHBP binds UCN and prevents it from activating CRHR1 (Seasholtz et al., 2002). Both overexpression of *Ucn* and KO of *Crhbp* improved RGC survival.

The second pair was *Timp2*, expressed by resRGCs and *Mmp12*, expressed by susRGCs. Administration of a soluble inhibitor of MMP12 was recently shown to improve RGC survival after ONC (Vinet et al., 2018), a result we replicated, and *Mmp12* deletion improves recovery from spinal cord injury (Wells et al., 2003). *In vivo*, MMP activity has been described to be closely controlled by endogenous tissue inhibitors (TIMP1-4) (Dzwonek et al., 2004) and AAV-mediated gene-transfer of *Timp1* and *Timp2* reduced neuronal damage in transient global ischemia, suggesting a neuroprotective role (Magnoni et al., 2007). The high expression of *Timp2* in resilient ipRGCs may locally block the proteolytic activity of MMPs, providing a survival advantage as their extracellular levels increase after ONC.

We additionally identified NDNF and PRPH as neuroprotective factors. In cultured hippocampal neurons, NDNF promotes neurite outgrowth and neuroprotection from glutamate and Aβ toxicity (Kuang et al., 2010). Certain isoforms of PRPH are cytoprotective against oxidative stress *in vitro* (McLean et al., 2014). To our knowledge, however, roles of NDNF and PRPH on neuron survival *in vivo* have not been reported.

A guiding hypothesis for this study was that protective genes identified on the basis of their expression by resRGCs would protect other RGC types if expressed broadly. We tested this hypothesis for *Ucn* and *Timp2*, and found that they improved the survival of CARTPT+ RGCs, which are among the most susceptible to ONC. This result supports the idea that targets identified by our methods could be broadly useful. On the other hand, these genes had no effect on survival of another susceptible subclass (NEUROD2+), indicating that neuroprotective strategies may need to be tailored to particular neuronal populations.

### Several neuroprotective targets also promote axon regeneration

Some of the targets identified in our screen for neuroprotection also promoted axon regeneration. Other interventions, such as depletion of *Pten*, have been shown to do both. However, these processes are not always linked and, in fact, can sometimes be antagonistic. For instance, deletion of *Dlk* promotes broad RGC survival but blocks Pten-mediated axon regeneration (Watkins et al., 2013), while *Sox11* overexpression promotes long distance axon regeneration in some RGCs but kills others (Norsworthy et al., 2017). Targets that do both may offer preferable therapeutic candidates in neurodegenerative contexts where axonal degeneration is implicated at early pathological stages.

### Distinguishing cell types from cell states

scRNAseq is increasingly used to generate cell atlases, in and outside of the nervous system (Regev et al., 2017). A general issue is whether some clusters of cells defined on the basis of transcriptional similarity represent different states of the same cell type, rather than different cell types. Because ONC leads to a dramatic but controlled change in cell state, our results provided an opportunity to explore this question.

Although injury-related gene expression changes were detectable at 0.5 and 1dpc, cells at this stage could be robustly assigned to type using a “one-step” classifier described above. On the other hand, classification at later times was increasingly impacted by stronger state-dependent changes. To reliably assign cells to types at late stages, we devised iGraphBoost, which iteratively assigns cells to types using a two-step approach that combines supervised classification and graph-based voting, while updating the model along the time course. We were able to map a high percentage of RGCs accurately even in highly degenerated retina. We expect that this approach will be effective when samples have sufficient temporal resolution to resolve gradual changes in molecular state from intrinsic programs.

In conclusion, we predicted and found that mining type-specific molecular correlates of resilience and vulnerability to injury provided a rich source of genes that mediate neuronal survival and axon regeneration. Some of the targets we found are likely to be effective in other contexts, and the general approach is likely to be applicable to other neuronal populations in the CNS.

## Acknowledgments

We thank Orenna Brand, Dustin Hermann, Allison Kao, Evan Macosko, Emily Marterskeck, Mu Qiao, Gevin Reynolds, Qingyang Wang and Yaxian Wang for assistance. This work was supported by grants from the NIH (MH105960, EY028625, EY021526, NS029169, NS104248, EY029360, EY028448, EY030204, EY026939, EY028625, P30EY012196 and AG056636), Wings for Life Spinal Cord Research Foundation and sponsored research agreements from Biogen Inc.

## Author Contributions

N.M.T., K.S., I.E.W., A.J., I.B., G.H., and J.R.S. conceived and designed experiments and analyzed data. N.M.T., I.E.W., A.J., I.B., G.H., W.C and M.E.A. performed experiments. K. S. developed the computational approaches. X.A. and J.Z.L. generated Smart-seq2 libraries. J.M.L and D.L. assisted with in vivo recordings. W.Y. analyzed data. C.M.L., A.R., Z.H. and J.R.S. provided supervision and acquired funding. N.M.T., K.S., I.E.W., A.J. and J.R.S. wrote the paper with input from all authors.

## SUPPLEMENTARY FIGURE LEGENDS

**Figure S1.**
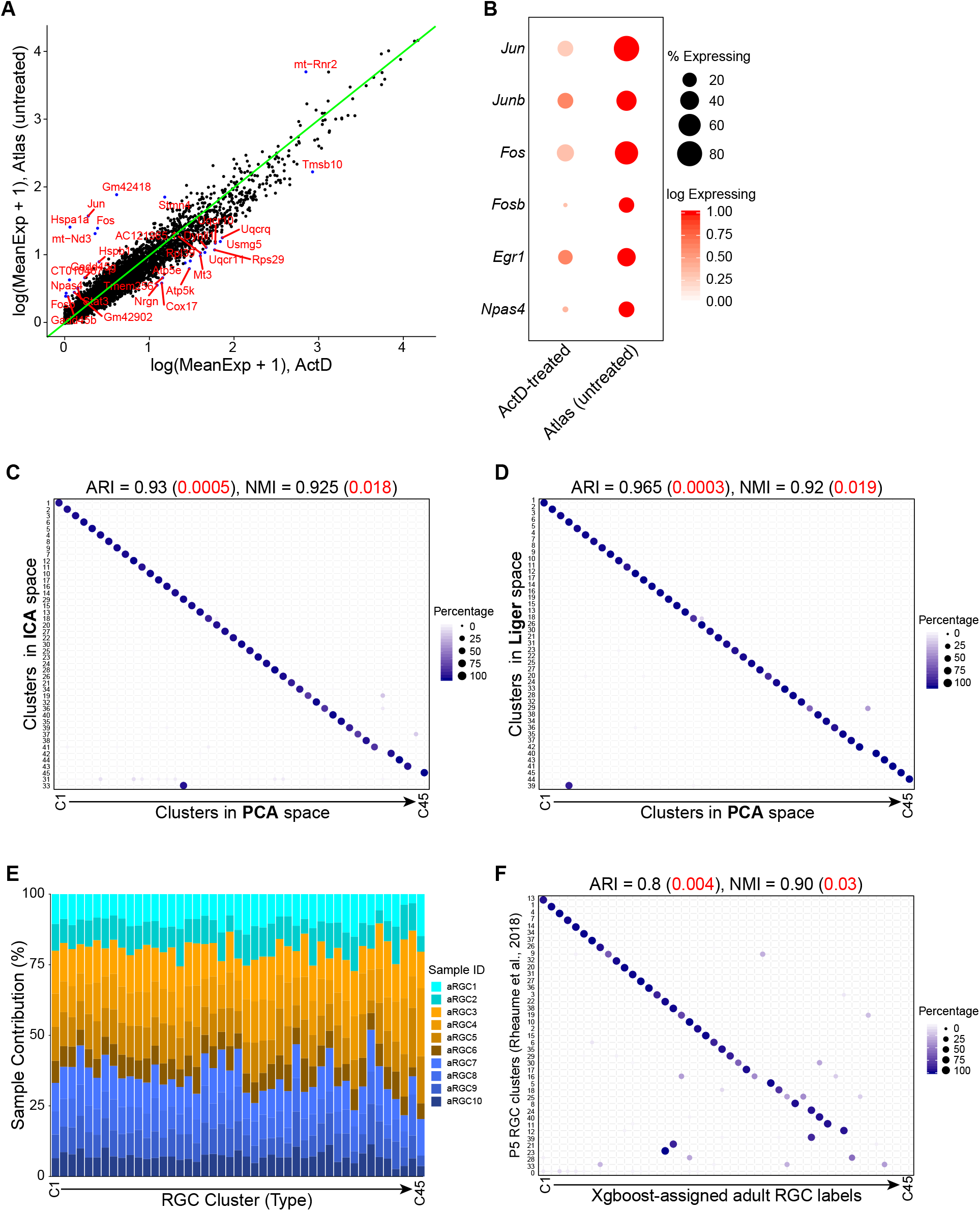
Assembling a molecular taxonomy of adult mouse RGC types, related to Figure 1. A. Scatter plot showing log_2_(mean expression+1) across genes in Act-treated 0dpc RGCs 0dpc (x-axis) and atlas RGCs (y-axis). Several DE genes (fold change > 2, p < 0.001, mAsT test) are highlighted (blue dots, red text). B. Dotplot showing lower expression of IEGs (rows) in Act-treated RGCs compared to atlas RGCs (columns). C. “Confusion Matrix” summarizing the correspondence between cluster labels determined using Louvain graph-clustering in a reduced dimensional space of gene expression computed using Principal Component Analysis (PCA, columns), versus a reduced dimensional space defined using Independent Component Analysis (ICA, rows). Circles and colors indicate the percentage of cells of a given row cluster that are members of a given column cluster. Agreement between data clusterings is quantified using two measures indicated on top – the adjusted rand index (ARI) and the normalized mutual information (NMI). ARI and NMI values range from 0 (random) to 1 (perfect match). Empirical ARI/NMI values suggested significant association compared to null ARI/NMI values from randomized associations indicated in parentheses (red color). D. Same as **C**, but rows indicate clusters computed in a reduced dimensional space computed by Liger (Welch et al., 2019), a non-negative matrix factorization (NMF) based approach. E. Stacked barplot showing the contributions of individual scRNA-seq samples (color) within each transcriptomically defined cluster (columns). For each cluster, the total contribution is normalized to 100%. F. Transcriptional correspondence between clusters of P5 RGCs (rows) reported by (Rheaume et al., 2018), and adult RGC types (columns) reported here. Circles and colors indicate the percentage of cells of a given P5 cluster that were assigned to an adult RGC type (column) based on a supervised, multi-class classification approach (see **Methods**).

**Figure S2.**
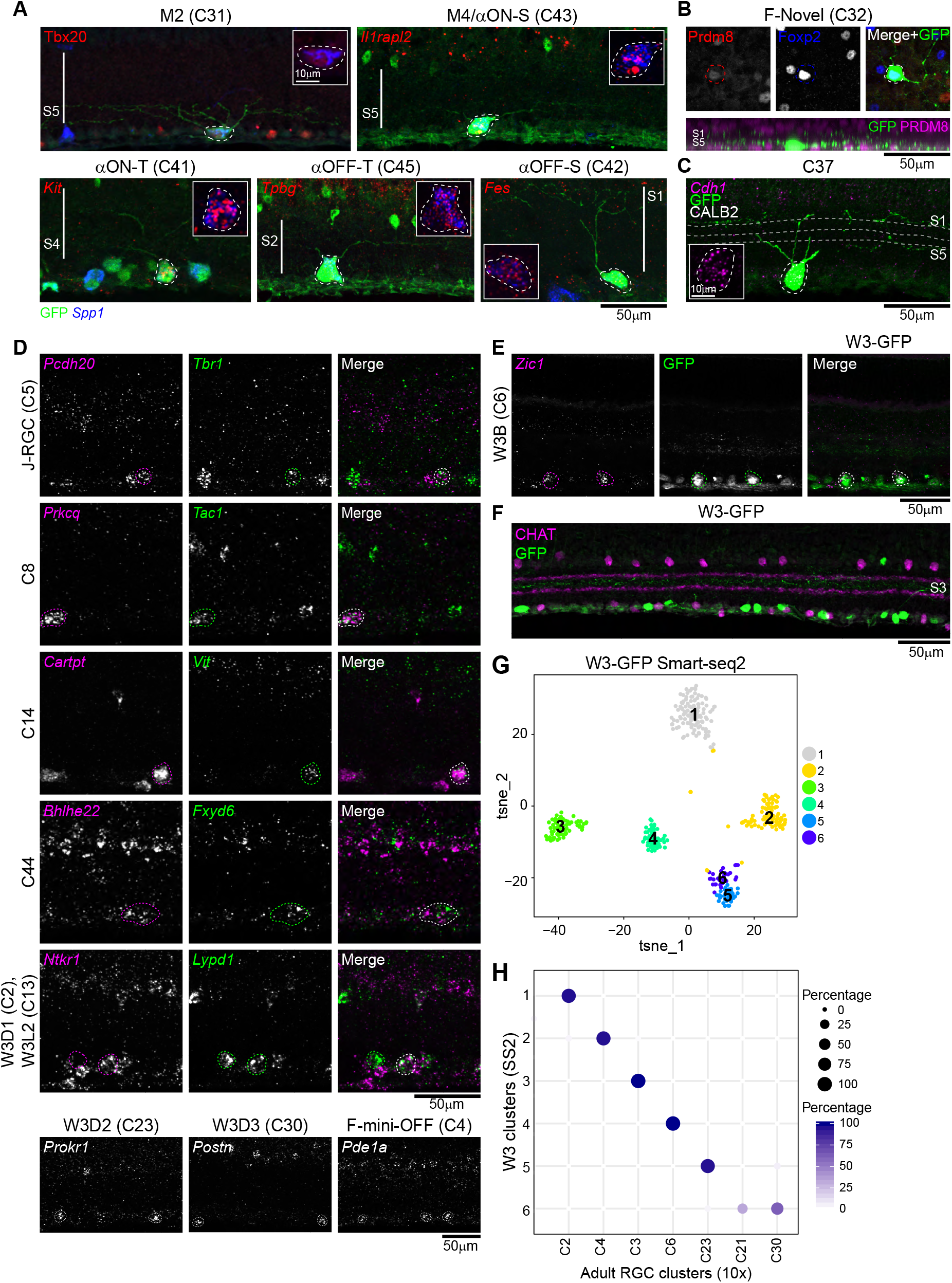
Histological characterization of molecularly defined RGC types, related to Figure 2. A. Combining FISH for *Spp1* (blue) and novel markers (red) listed in **Figure 2B** with IHC shows that RGCs (sparsely labeled in YFP-H line, green) include the 4 αRGC types, and the ipRGC type known as M2. B. C32 (F-Novel) is an S1/5 bistratified RGC (labeled in green) that co-expresses FOXP2 (blue) and PRDM8 (red). Cross-section generated from rotation of *en face* wholemount image. C. Novel RGC type C37 laminates just above and below S2 and S4, respectively. Shown by combining FISH for *Cdh1* (magenta) and IHC on sparsely labeled RGCs in the YFP-H line (green). IPL sublaminae, drawn from CALB2 staining are indicated (white). D. Validation of molecular signatures of 9 RGC types in the atlas using 1 and 2-color FISH. Included are (from top to bottom), J-RGC/C5 *(Tbr1+Pcdh20+)*, C8 *(Tac1+Prkcq+)*, C14 *(Cartpt+Vit+)*, C44 *(Bhlhe22+Fxyd6+)*, W3D1/C1 *(Lypd1+Ntrk1-)*, W3L2/C13 *(Lypd1 +Ntrk1* +), W3D2/C23 *(Prokr1+)*, W3D3/C30 *(Postn+)* and F-mini-OFF/C4 *(Pde1a)*. E. Validation of *Zic1* expression in W3B (C6) by combining FISH (magenta) plus IHC on the *W3-GFP* line (green). F. The W3-GFP mouse line (green) labels multiple RGC types that laminate predominantly in S3, seen here between the S2/4 sublaminae of the IPL labeled using an antibody for CHAT (magenta) G. t-SNE visualization of RGCs from the W3-GFP line, sequenced using the Smart-seq2 protocol (Picelli et al., 2013). Numbers indicate clusters identified by graph clustering. H. Supervised classification analysis shows that 5/6 W3-GFP RGC clusters (rows) map 1:1 to 5 clusters in the adult RGC atlas (columns), with the remaining cluster mapping 1:2.

**Figure S3.**
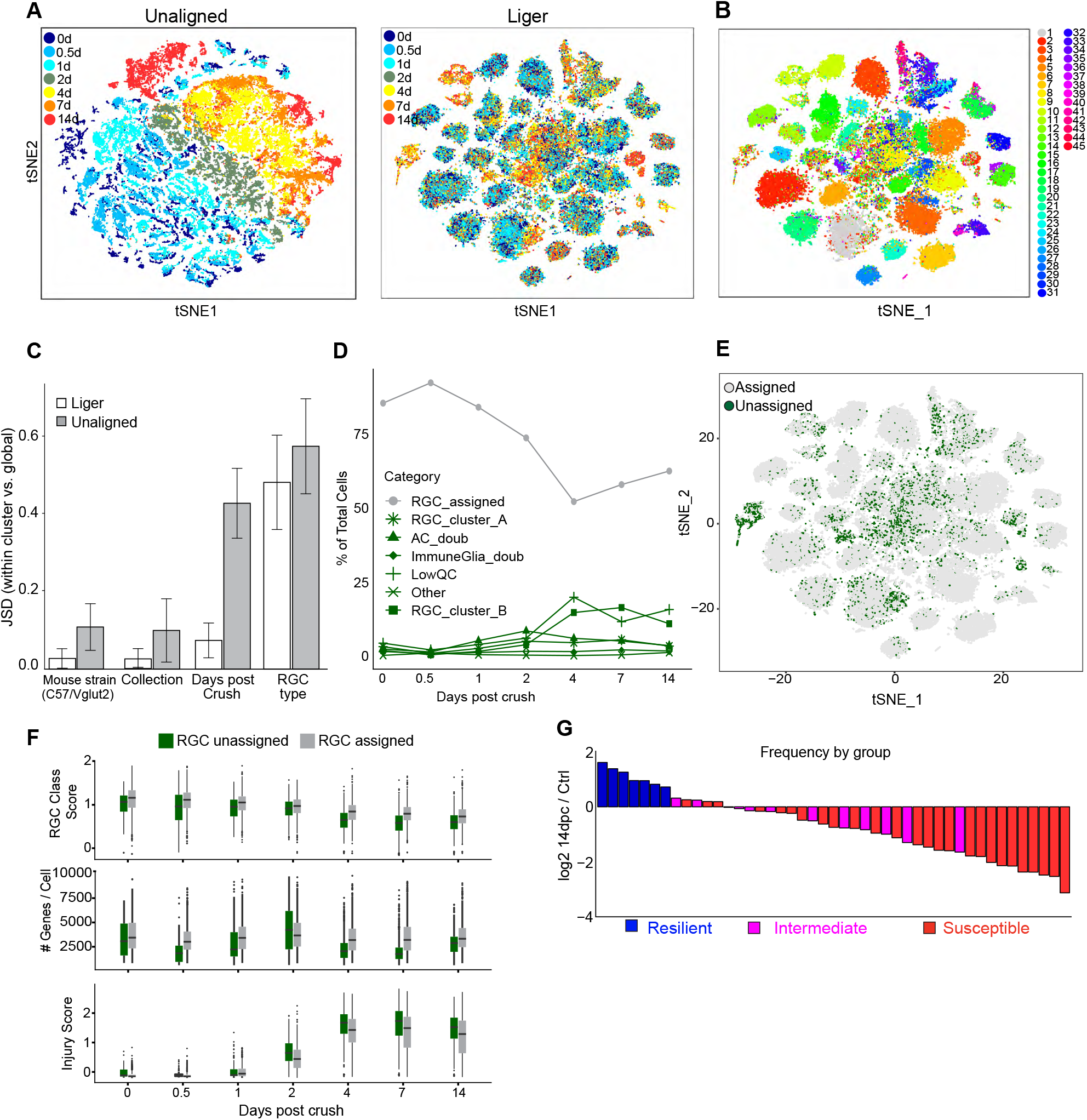
iGraphBoost accurately estimates survival of RGC types following ONC, related to Figure 3. A. t-SNE visualization of 76,646 post-ONC (including 0dpc) RGCs based on PCA without temporal alignment (left), and Liger (right). Cells are colored by time points (colored dots). B. t-SNE visualization of injured RGCs colored by iGraphboost type assignments. Unassigned RGCs are not plotted. Note that well-separated groups in t-SNE, a 2D visualization tool that does not fully capture features in higher-dimensions, are predominantly (but not entirely) comprised of individual types. C. The influence of various cell metadata (x-axis) on clusters identified using PCA (without alignment, grey) and Liger (white). Bars show the Jensen Shannon Divergence (JSD) computed between the metadata composition within a cluster and the global background. mean ±SD computed across clusters. While clusters defined using the Louvain-Jaccard approach in both PCA and Liger spaces are enriched for specific RGC types, Liger-based clusters are significantly less influenced by time post injury compared to PCA clusters, suggesting a better alignment of RGC types along the injury time course. D. Frequency of transcriptomically-defined subgroups within unassigned RGCs along the time course. Two unassigned subgroups appear to be doublets (amacrine+RGC+ (AC_doub), immune cells/glia+RGC+ (ImmuneGlia_doub) based on the co-expression of RGC and non-RGC markers (**Figure 1E**). Two groups showed expression of RGC markers but were not enriched with sufficient type-specific markers to be assigned (RGC_cluster_A and B), one of which (RGC_cluster_B) increased in frequency at 4-14dpc. Two groups were comprised of low quality cells based on numbers of genes and transcripts recovered (Other, LowQC). E. t-SNE visualization (as in **Figure S3A**, *right panel* (Liger)) of injured RGCs that are assigned (grey dots) or unassigned (green dots) to atlas types by iGraphboost. Unassigned cells were distributed throughout Liger clusters but were most concentrated in clusters composed primarily of 4-14dpc RGCs (see panel A), suggesting that the injury-related state changes at these later time points prevented their classification by iGraphboost. F. Box and whisker plots show differences in key metrics (y-axis) between RGCs that passed initial quality metrics that could and could not be assigned (colors) (groups RGC_unassigned and RGC_unassigned_late from panel E) as a function of time (x-axis) post ONC. On average, unassigned RGCs show lower expression of RGC class genes (top), lower number of genes per cell (middle), and higher expression of injury response genes (bottom), particularly from 4-14dpc. Black horizontal line, median; bars, interquartile range; vertical lines, minimum and maximum; dots, outliers. G. RGC types ranked by 14dpc survival (as in **Figure 3E**) colored by survival group (**Figures 3J-L**).

**Figure S4.**
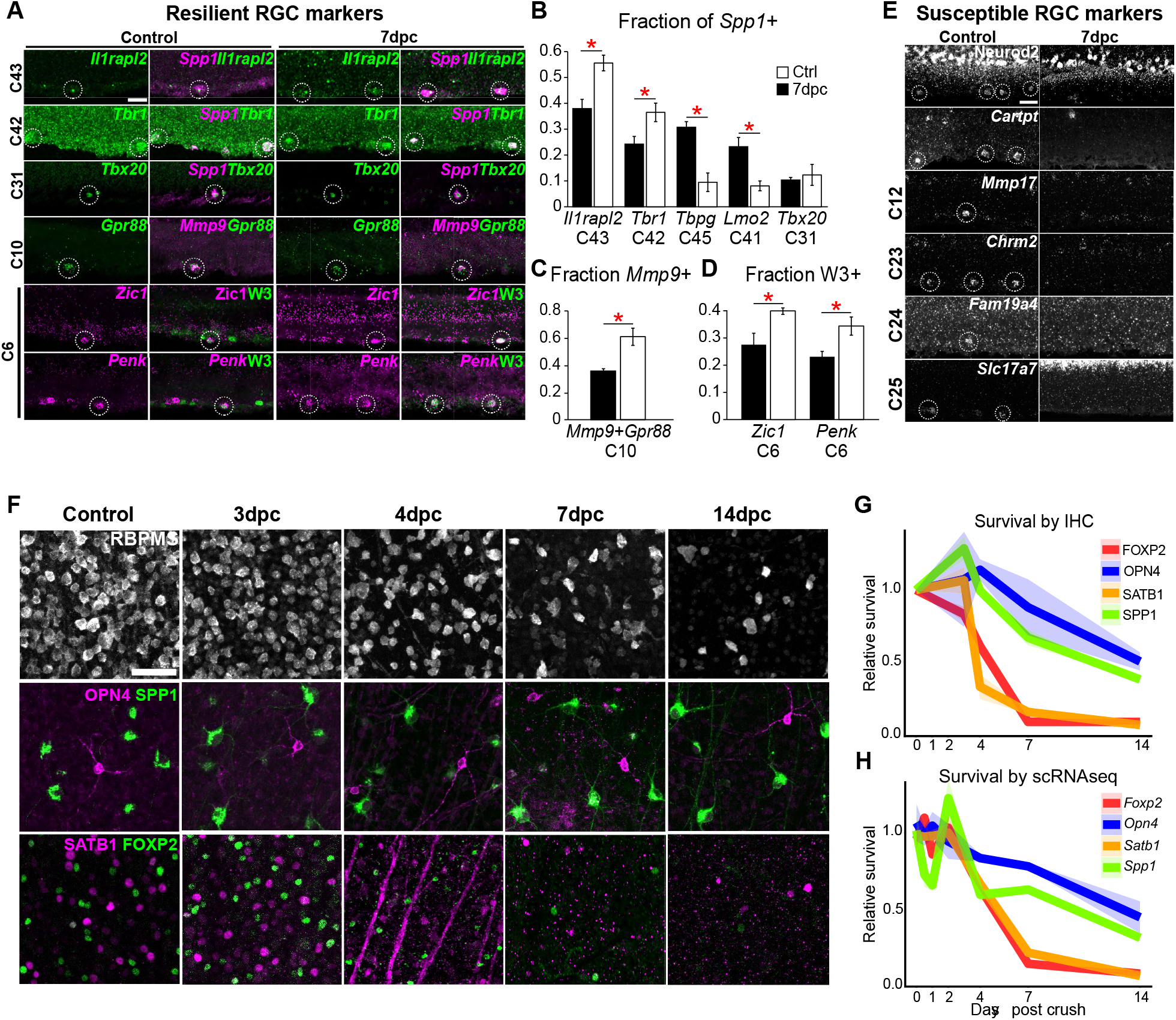
Immunohistochemical validation of scRNA-seq-derived survival rates, related to Figure 3. A. Identification of 4 resRGC types (C10, 31, 42, 43), and an intRGC type (C6) using a combination of FISH (italicized) and/or IHC (capitalized). Shown are 6 staining combinations that individually target these types at control and 7dpc. ‘W3’ indicates IHC for GFP in W3-GFP mouse line. White circles indicate double labeled cells. Scale bar: 25μm (Markers detailed in **Figure 2**) B. Quantification of FISH/IHC from **A**. Comparison of the relative proportions of five SPP1 + RGC types at 0 and 7dpc shows that the relative proportions of resRGC types (C43: *Il1rapl2+* and C42: *Tbr1+)* increase, while those of susRGC types (C45: *Tpbg)* and (C41: *Lmo2)* decrease. resRGC (C31: *Tbx20)*, which has lower 14dpc survival than C43 and C42 by scRNA-seq, did not change in proportion significantly. Results are consistent with scRNA-seq rankings. (*p<0.05, Student’s t-test, error bars: SEM). Note that by IHC marker quantification, C43 (αRGc ON-S/M4) was the most abundant, while C31 (ipRGC M2) was the least, in contrast to their measured frequency by scRNA-Seq (**Figure 1D**), suggesting a possible sampling bias in the latter. (n≥4 retinal sections from at least 2 mice used for each quantification) C. Quantification of FISH/IHC from **A**. Fraction of *Mmp9+* RGCs that are also *Gpr88+* (C10) at 0 and 7dpc. Increase in relative proportion of C10 at 7dpc highlights its relative resilience, consistent with scRNA-seq ranking. (*p<0.05, Student’s t-test, error bars: SEM). D. Quantification of FISH/IHC from **A**. Fraction of W3+ (GFP labeling of *W3* mouse line) RGCs labeled by *Zic1* or *Penk*, which label C10 (W3B) at 0 and 7dpc. Quantifications show that W3B RGCs, an *intRGC*, has a higher relative resilience than other W3 types, consistent with panel **B** and scRNA-seq rankings. (*p<0.05, Student’s t-test, error bars: SEM). E. FISH (italicized) or IHC (capitalized) frequencies for markers labeling different *susRGC* populations (markers detailed in **Figure 2**). In each case, the signal is strongly reduced at 7dpc indicating RGC loss (right column). Scale bar: 25μm. F. Whole mount IHC micrographs showing RGC loss along ONC time course for a pan-RGC marker RBPMS (top row), and for 4 RGC subclass markers including OPN4 (middle row; labels resRGC types C33, 40), SPP1 (middle row), SATB1 (bottom row) and FOXP2 (bottom row). SPP1, SATB1, FOXP2 labeled clusters as in **A**. For Scale bar: 50μm. G. IHC based quantification of survival (density at each timepoint / density in control) for RGC subclasses labeled by markers in **F**. Color ribbons: +/− SEM. H. scRNA-seq-based quantification of survival for RGC subclasses labeled by markers in panel **F** demonstrates striking agreement with IHC trends in panel **F**. Color ribbons: +/− SD.

**Figure S5.**
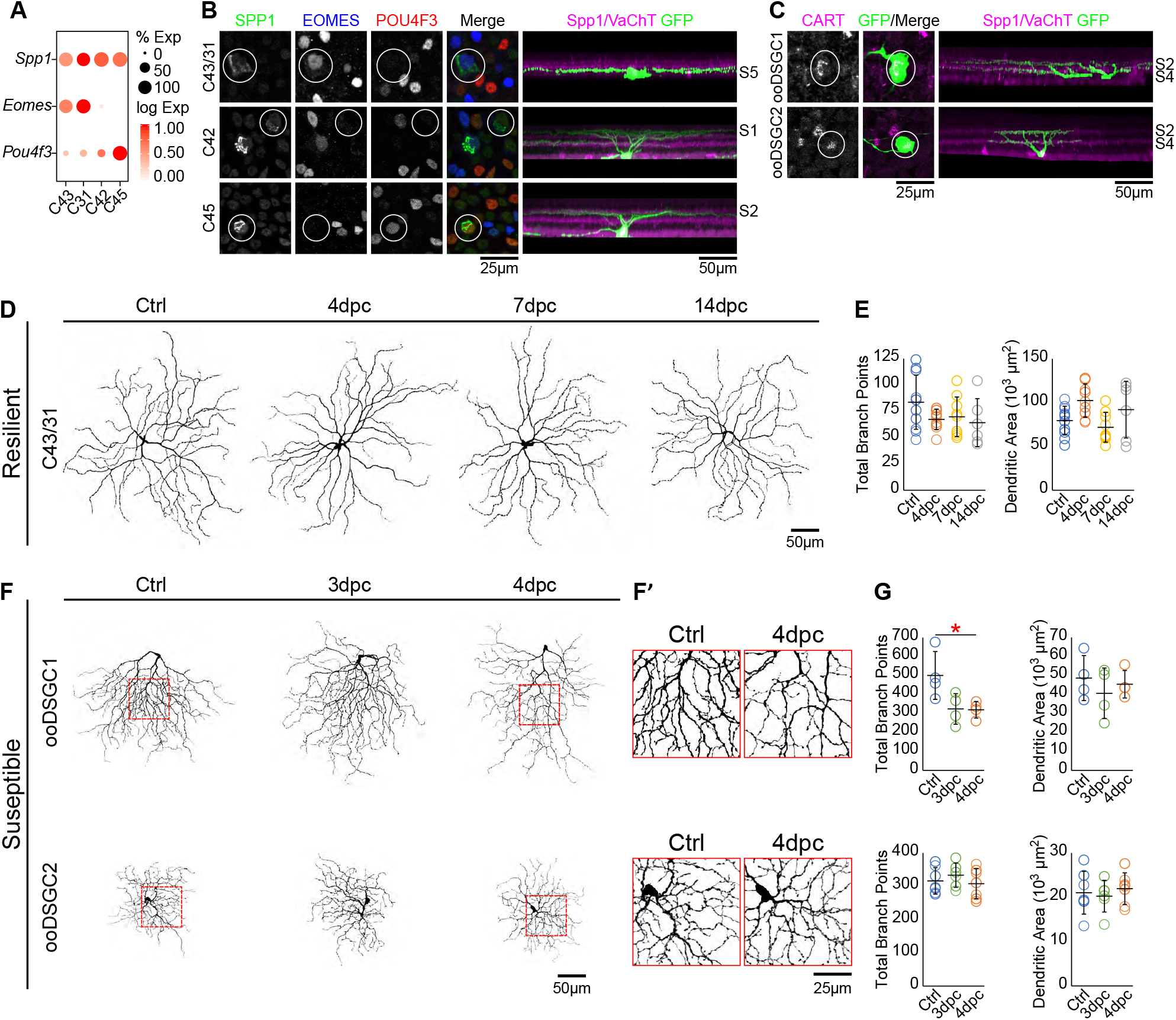
Morphological changes in RGCs following ONC. A. Dotplot showing expression patterns of markers used to label *Spp1+* RGC types in **Figure 5M** and panel **D**. B. IHC for markers in **A** on whole-mount YFP-H retinas labels (top to bottom) αRGC/ipRGC ON-S (C43) and ipRGC M2 (C31) (both SPP1 +EOMES+POU4F3-), αRGC OFF-S (C42) (SPP1 +EOMES-POU4F3-), and αRGC OFF-T (C45) (SPP1 +EOMES-POU4F3+). *YFP-H* labeling (green) in rotational sections illustrates distinct dendritic lamination patterns of these SPP1+ RGC types. C. IHC for CART and GFP on whole-mount YFP-H retinas labels S2/S4 laminating ooDSGCs, here showing examples for the asymmetric *Hb9*-like and small/bushy ooDSGCs D. Example *en face* morphology of an additional resilient type (identified either as ipRGC M2 (C31) or αON-S (C43)) at Ctrl, 4, 7, and 14dpc. E. Quantification of C31/42 morphological complexity (total branch points, left) and size (dendritic area, right) shows no significant difference between time points for either measure (one-way ANOVA with post-hoc Tukey HSD test). Data are shown as mean ±SD. F. Example *en face* morphology of two distinct susceptible types ooDSGC1 and ooDSGC2 (*Hb9*-like and small/bushy, respectively) at Ctrl, 3, and 4dpc. **F’** showing zoomed in views of dendrites at Ctrl and 4dpc. G. Quantification of ooDSGC1 morphological complexity (total branch points, left) shows a significant difference between Ctrl and 4dpc (* = p<0.05) while ooDSGC2 does not. Neither ooDSGC type shows no significant difference of size (dendritic area, right) between time points (one-way ANOVA with post-hoc Tukey HSD test). Data are shown as mean±SD. ooDSGC1 n=4, 4, 4 cells for Ctrl, 3, and 4dpc, respectively.

**Figure S6.**
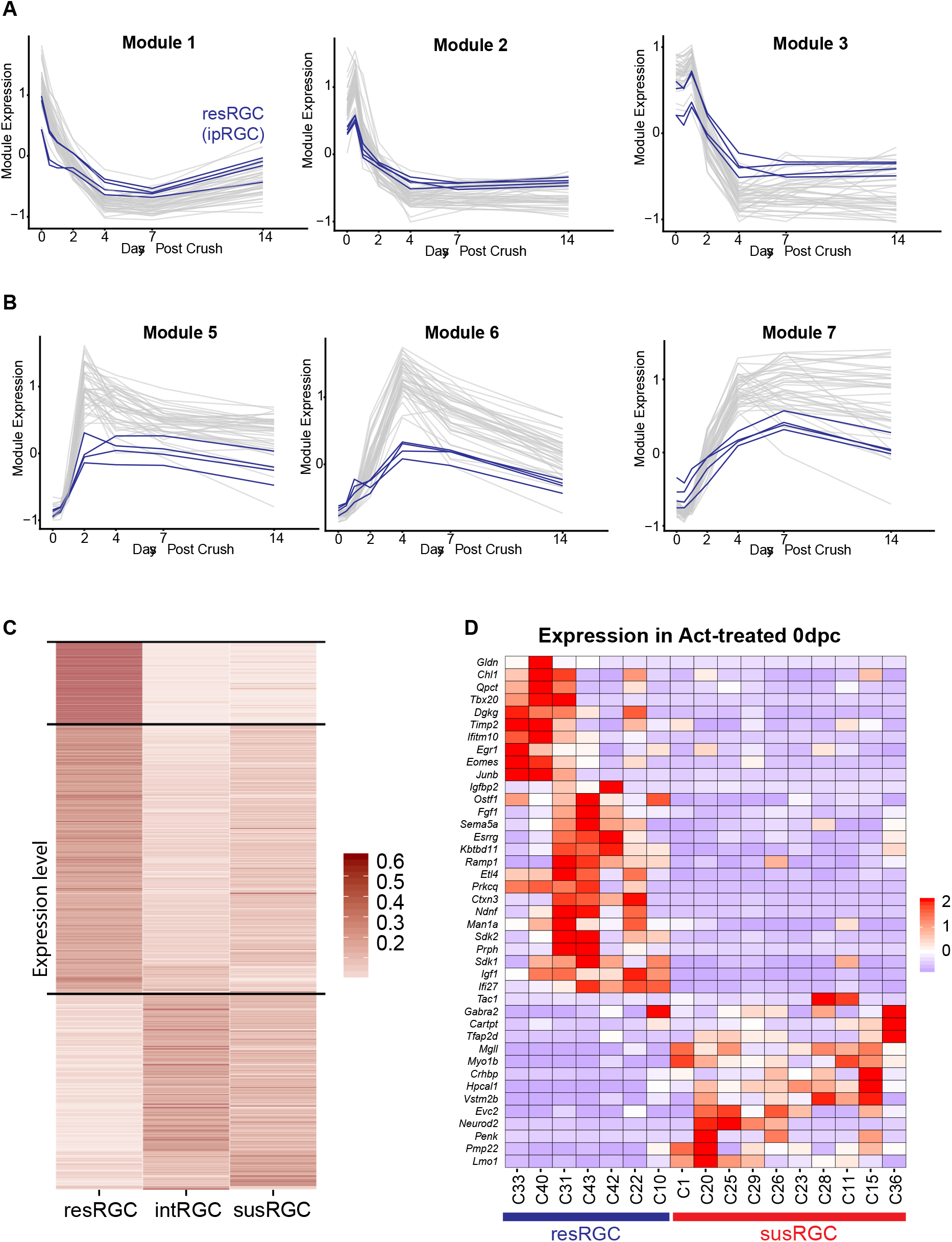
Pre and post-injury gene expression changes in RGCs and effects of actinomycinD treatment, related to Figure 6. A. Temporal dynamics of the average expression of globally-identified downregulated gene Modules 1, 2, and 3 in **Figure 5A**. Genes plotted by cluster (lines). Blue lines correspond to 4/5 ipRGC types (C31, 22, 40, 33). We did not observe strong differences in the degree of downregulation among RGC types. B. As in **A**, but for gene Modules 5, 6, and 7. ipRGCs (blue lines) showed consistently weaker average upregulation of genes in these modules. C. Heatmap of the averaged transcript level of differentially expressed genes between survival groups (*res, int*, and *susRGCs*) in control RGCs. D. Same as **Figure 5F**, but calculated from Act-treated RGC transcriptomes.

**Figure S7.**
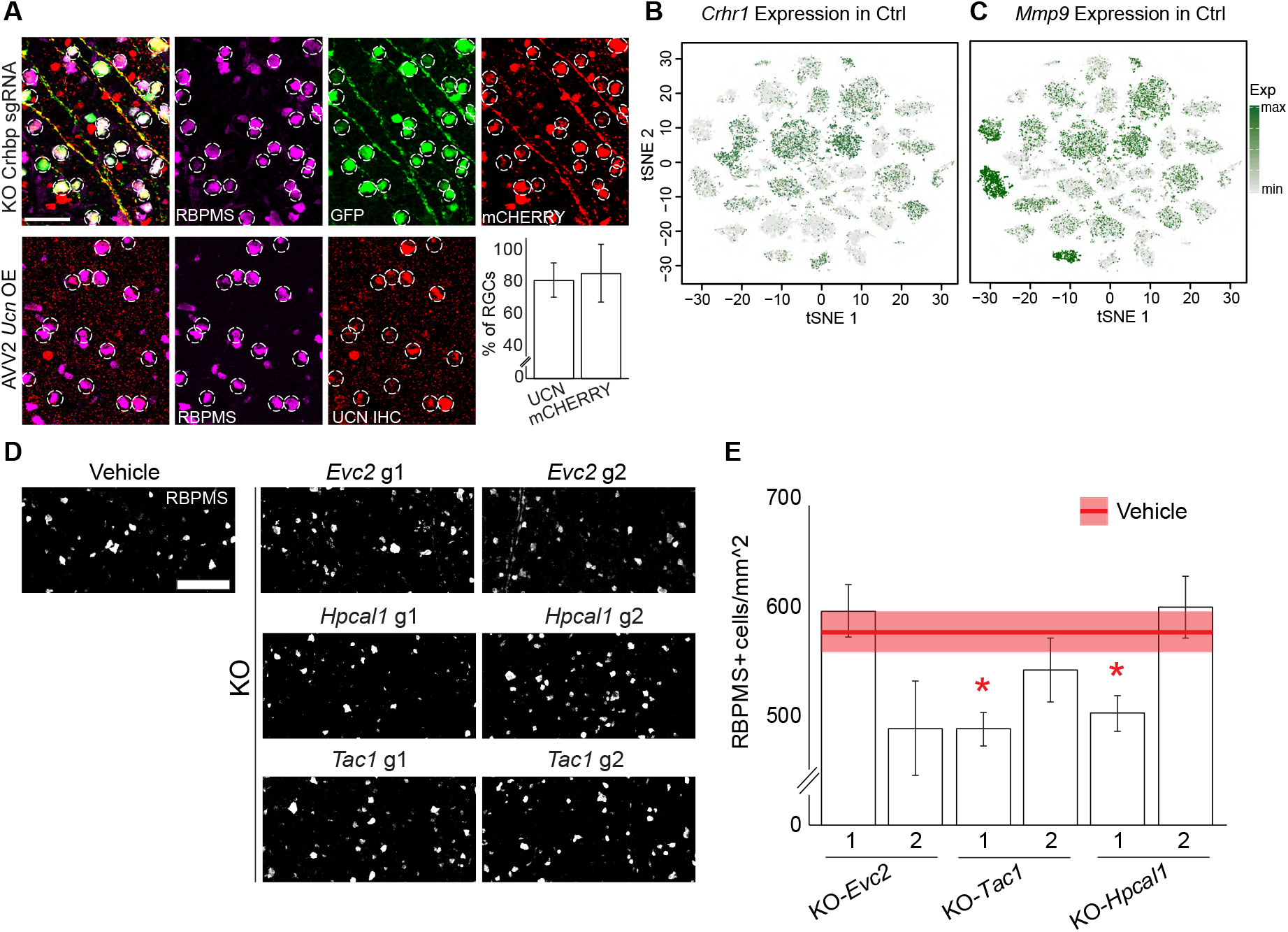
in vivo testing of genes associated with resilience or vulnerability, related to Figure 7. A. Transfection efficiency of AAV2 demonstrated by AAV2 mediated delivery of mCHERRY tagged *Crhbp* sgRNA#2 in *Vglut2-Cre* x *LSL-Cas9-Gfp* mice (upper) or UCN via IHC in *C57Bl6/J* mice (lower) (Scale bar: 50μm). White circles show RGCs positive for RBPMS, GFP and mCHERRY; or RBPMS and UCN. *Lower right:* Quantification shows % of RBPMS^+^UCN^+^ (n=2) or RBPMS^+^mCHERRY^+^ *(Crhbp* sgRNA #2) (n=7) double positive RGCs (error bars = SD). B. *Crhr1* expression in multiple Atlas RGC types visualized by tSNE, as in **Figure 1C**. Cells are colored based on their expression level in a monochrome scale, with darker green indicating higher expression. C. Same as panel B, for *Mmp9* expression. D. *En face* whole mount images of IHC for RBPMS following the indicated intervention at 14dpc. E. IHC quantification of RBPMS density (RGCs/mm^2^) in retinal whole mounts following the indicated interventions (KO-*Evc2, Tac1* or *Hpcal1*), no positive effect on survival was observed for any sgRNA. *adjusted p-value <.05 (FDR).

## Supplementary Table Legend

**Table S1.** Overview of molecularly defined RGC clusters, cross-referenced to types previously defined based morphology, function and molecular signatures

| Type | This paper | Bae et al. 2018 | Baden et al., 2016 | Description of function<sup>a</sup> |
| --- | --- | --- | --- | --- |
|  |  | 1ni | G3-OFF step | OFF response (Bae et al. 2018) |
|  |  | 1no | G3-OFF step | OFF response (Bae et al. 2018) |
| M1 | 33 | 1ws |  | ipRGC, weak light evoked response (Bae et al 2018) |
| alpha OFF S | 42 | 1wt | G5a,b-OFF S alpha | sustained OFF response, NSGC (Krieger et al. 2017) |
| F-mini OFF | 4 | 2an |  | transient OFF response, DSGC (Rousso et al. 2016) |
| F-midi OFF and J-RGCs | 28,5 | 2aw | G4,6-OFF slow,(ON-)OFF Jam-B mix | F-midi OFF: sustained OFF, NSGC (Rousso et al. 2016);<br>J-RGC: OFF, DSGC, color-opponent (Kim et al. 2008, Joesch & Meister 2016) |
|  |  | 2i |  | OFF response (Bae et al. 2018) |
|  |  | 2o | G5b,c-OFF S alpha-like | OFF response (Bae et al. 2018) |
|  |  | 25 |  |  |
| Cdh1-RGC | 37 | 27?<sup>b</sup> |  |  |
|  |  | 28 |  |  |
| Tbr1-S1 | 17 | 3i? |  |  |
|  |  | 3o |  | OFF response (Bae et al. 2018) |
| ooDSGC, posterior (nasal) | 12 | 37c | G12,13-ON-OFF DS | transient ON-OFF response, DSGC (Vaney, Sivyer, Taylor 2012) |
| ooDSGC, superior (dorsal) | 16 | 37d | G12,13-ON-OFF DS | transient ON-OFF response, DSGC (Vaney, Sivyer, Taylor 2012) |
| ooDSGC, anterior (temporal) | C10 or C24 | 37r? | G12,13-ON-OFF DS | transient ON-OFF response, DSGC (Vaney, Sivyer, Taylor 2012) |
| ooDSGC, inferior (ventral) | 16 | 37v | G12,13-ON-OFF DS | transient ON-OFF response, DSGC (Vaney, Sivyer, Taylor 2012) |
| Tbr1-S2 | 21 | 4i? | G9-OFF T alpha mini |  |
|  |  | 4on | G9-OFF T alpha mini | OFF response (Bae et al. 2018) |
| alpha OFF T | 45 | 4ow | G8-OFF T alpha | transient OFF response, NSGC (Krieger et al. 2017) |
|  |  | 5to |  |  |
| HD1 or 2 Jacoby and Schwartz |  | 5si | could be G11-ON-OFF-local |  |
| HD1 or 2 Jacoby and Schwartz |  | 5so | could be G11-ON-OFF-local |  |
| UHD Jacoby and Schwartz |  | 5ti | could be G11-ON-OFF-local |  |
| W3B/LED | 6 | 51 | G10-ON-OFF local-edge | transient response, object motion (Zhang et al. 2012) |
| F-mini ON | 3 | 63 |  | transient ON response, DSGC (Rousso et al. 2016) |
|  |  | 6sn | G23-ON alpha mini | ON response (Bae et al. 2018) |
| alpha ON T | 41 | 6sw | G19-ON T alpha (lam mismatch) | transient ON response, NSGC (Krieger et al. 2017) |
| F-midi ON | 38 | 6t |  | transient ON response, NSGC (Rousso et al. 2016) |
| ON DS S |  | 7id | G25,26,29-ON DS S | sustained ON response, DSGC (Bae et al. 2018) |
| ON DS S |  | 7ir | G25,26,29-ON DS S | sustained ON response, DSGC (Bae et al. 2018) |
| ON DS S |  | 7iv | G25,26,29-ON DS S | sustained ON response, DSGC (Bae et al. 2018) |
| ON DS T |  | 7o | G16-ON DS T | transient ON response, DSGC (Bae et al. 2018) |
|  |  | 72 |  | sustained ON response (Bae et al. 2018) |
| ON-delayed |  | 73 |  | sustained ON response (Bae et al. 2018) |
| Vglut1-RGC | 25 | 8n? |  | ON response (Bae et al. 2018) |
| alpha ON S | 43 | 8w | G24-ON alpha | ipRGC, sustained ON response, NSGC (Krieger et al. 2017) |
| F-novel | 32 | 81i? |  | ON response (Bae et al. 2018) |
|  |  | 81o |  |  |
|  |  | 82n |  | ON response (Bae et al. 2018) |
| vertical orientation selective cell |  | 82wi |  | sustained ON response (Bae et al. 2018) |
|  |  | 82wo |  |  |
|  |  | 85 |  | ON response (Bae et al. 2018) |
|  |  | 9n |  | sustained ON response (Bae et al. 2018) |
| M2 | 31 | 9w |  | ipRGC, sustained ON response, NSGC (Bae et al. 2018) |
|  |  | 91 |  | ON response (Bae et al. 2018) |
|  |  | 915 |  |  |

<sup>a</sup>orientation selective (OSGC), direction-selective (DSGC), non-selective (NSGC), intrinsically photosensitive (ipRGC)
<sup>b</sup>Uncertain type pairings are indicated with a '?'

**Table S2.**
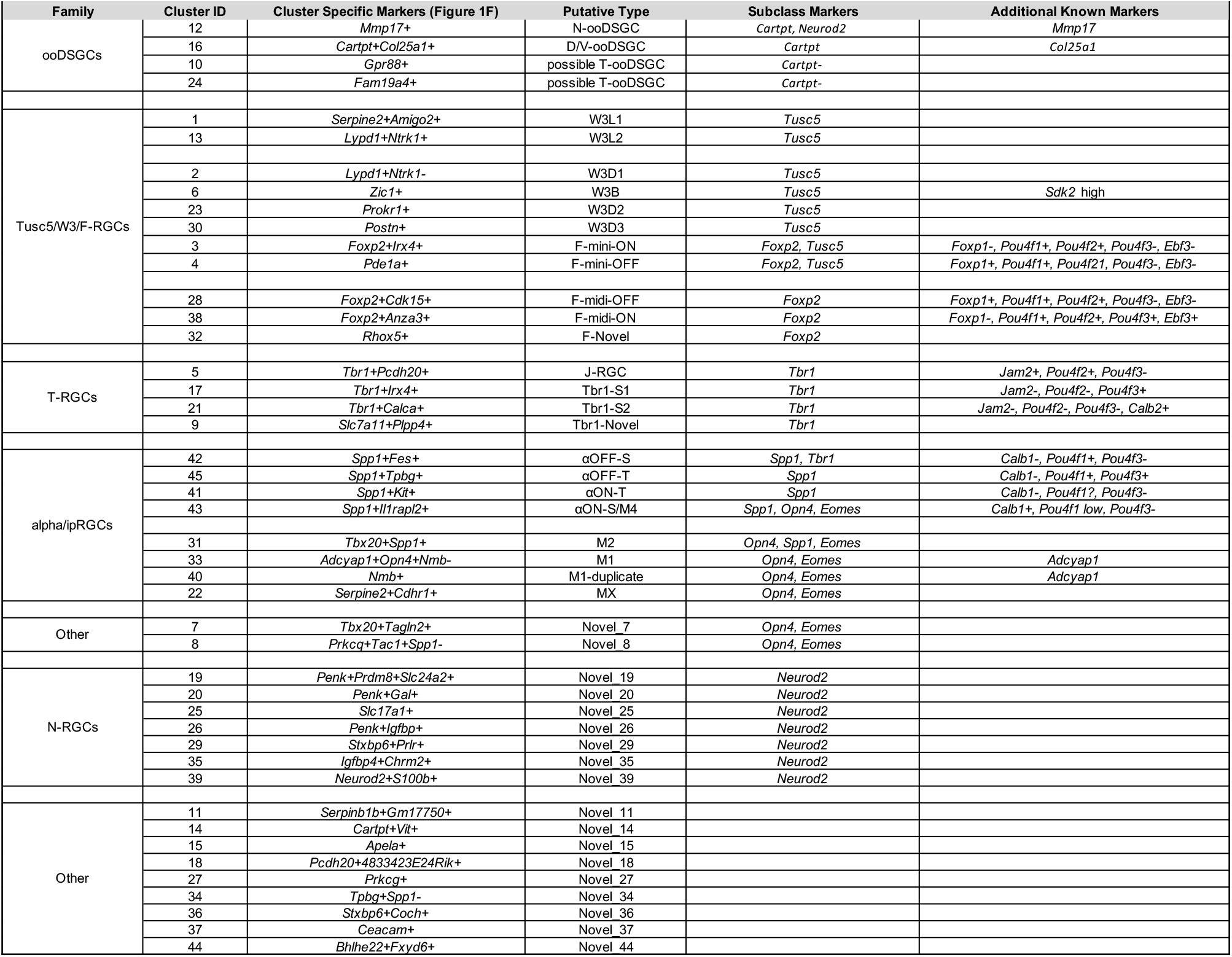
Established molecular markers for RGC subclasses and types and explanation of naming convention for clusters

**Table S3.** RGC subclass survival estimates by scRNAseq and IHC at 14dpc. Summary of combinatorial marker RGC subclass survival as calculated by scRNAseq compared to retinal whole mount IHC staining quantifications at 14dpc.

| Antibody and/or mouse line | Clusters labeled | Subgroup survival<br>scRNA-Seq | Subgroup survival<br>IHC | IHC<br>SEM | n | p value<sup>b</sup><br>(Stain<br>vs. RBPMS) |
| --- | --- | --- | --- | --- | --- | --- |
| 1 SPP1+; *Opn4-CreXROSA TdTomato* | 31, 43<sup>a</sup> | 0.853 | 0.889 | 0.147 | 8 | <0.001* |
| 2 OPN4+ | 33, 40 | 0.534 | 0.564 | 0.102 | 6 | 0.002* |
| 3 SMI32 | 42, 43, 45 | 0.537 | 0.526 | 0.108 | 5 | 0.004* |
| 4 *Opn4-CreXROSA TdTomato* | 22, 31, 33, 40, 43 | 0.478 | 0.512 | 0.025 | 8 | <0.001* |
| 5 *Opn4-CreXROSA TdTomato*; SPP1-; OPN4- | 22 | 0.452 | 0.464 | 0.059 | 3 | 0.010* |
| 6 SPP1 | 41, 42, 43, 45 | 0.377 | 0.381 | 0.037 | 15 | <0.001* |
| 7 *Cck-CreXYfp STP15*; SPP1+ | 31<sup>a</sup> | 0.361 | 0.314 | 0.019 | 3 | 0.010* |
| 8 W3 (bright cells) | 6 | 0.244 | 0.286 | 0.066 | 3 | 0.186 |
| 9 W3, FOXP2- | 2, 6, 23 | 0.201 | 0.256 | 0.041 | 3 | 0.243 |
| 10 SPP1+; *Opn4-CreXROSA TdTomato*- | 41, 42, 45 | 0.228 | 0.243 | 0.040 | 9 | 0.305 |
| 11 W7; SPP1+ | 42, 45 | 0.337 | 0.238 | 0.025 | 6 | 0.159 |
| 12 *Vglut2-CreXYfp STP15* | All RGC's | ND | 0.209 | 0.019 | 6 | 0.315 |
| 13 RBPMS | All RGC's | ND | 0.188 | 0.016 | 11 |  |
| 14 W3 | 2, 3, 4, 6, 23 | 0.174 | 0.181 | 0.030 | 3 | 0.815 |
| 15 *Pv-CreXYfp STP15* | PV+ RGC's | 0.096 | 0.135 | 0.022 | 4 | 0.117 |
| 16 *Cck-CreXYfp STP15* | Cck+ RGC's | 0.306 | 0.132 | 0.011 | 5 | 0.047* |
| 17 W3 (Dim cells) | 2, 3, 4, 23 | 0.159 | 0.115 | 0.008 | 3 | 0.036* |
| 18 *Cck-CreXYfp STP15*; SATB1+ | 14, 16 | 0.085 | 0.096 | 0.028 | 4 | 0.019* |
| 19 W3, FOXP2+ | 3, 4 | 0.140 | 0.093 | 0.042 | 3 | 0.036* |
| 20 *Jam-CreerXYfp STP15* | 5 | 0.152 | 0.087 | 0.005 | 3 | 0.010* |
| 21 FOXP2 | 3, 4, 28, 32, 38 | 0.124 | 0.083 | 0.018 | 6 | 0.003* |
| 22 *Pv+ RGC's*; SPP1- | non-α PV+ RGC's | 0.065 | 0.077 | 0.007 | 4 | 0.004* |
| 23 *Hb9* | 16 | 0.110 | 0.064 | 0.015 | 3 | 0.010* |
| 24 SATB1 | 6, 12, 13, 15, 16, 26, 34-36 | 0.106 | 0.062 | 0.025 | 4 | 0.009* |
| 25 SATB2 | 9, 12, 19, 20, 25, 26, 29, 35, 39 | 0.065 | 0.037 | 0.016 | 3 | 0.010* |

<sup>a</sup>For staining combination #1, we report the scRNAseq survival estimate for C43 alone, while this IHC combination labels both C31 and C43. This was necessitated by a small sampling bias against Alpha RGCs compared to ipRGCs in scRNAseq (see *Spp1* (Alpha RGC marker) vs *Eomes* (ipRGC marker) in **Figure 1G** and direct comparison of C31, C43 in **Figure S4B**), which distorted their combined assessment. A more accurate measure for C31 survival is given in staining combination #7.
<sup>b</sup>For IHC error measurements: \*p<.05, Kruskal-Wallis rank order test

**Table S4.**
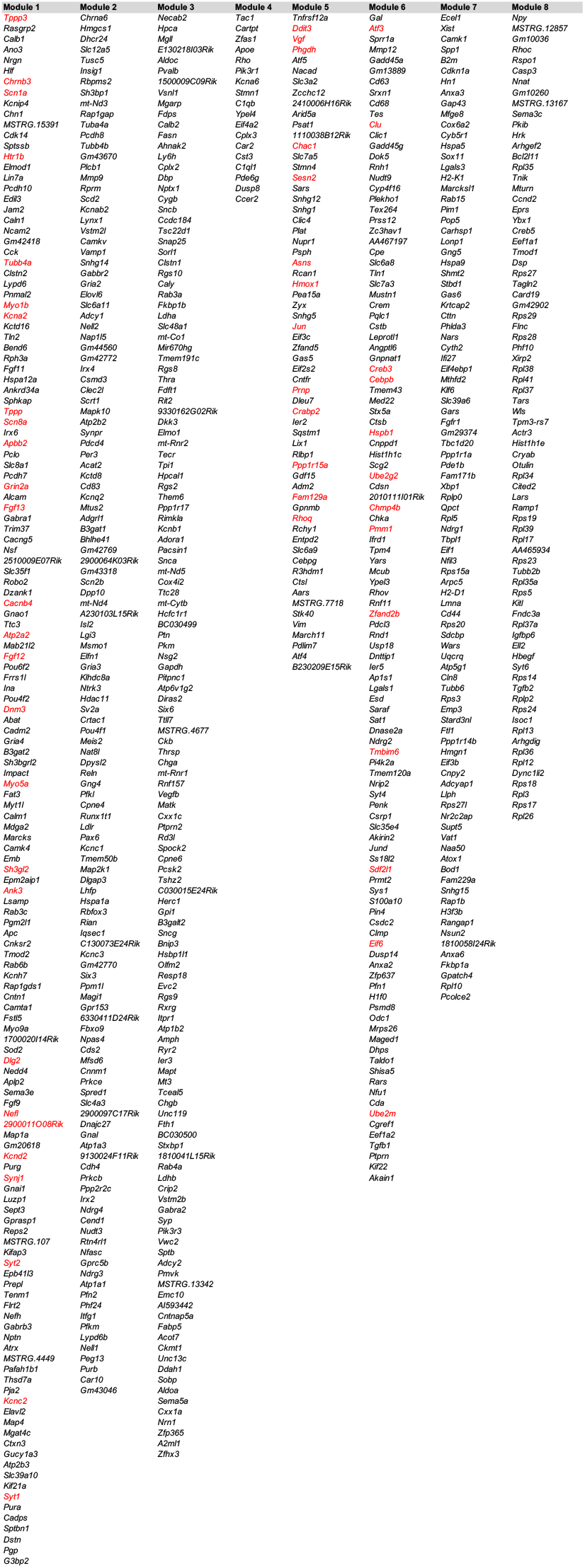
Globally regulated gene modules (Mod1-8) over ONC time course. Red text indicates Module 1 genes plotted in Figure 5B or Module 5,6 genes plotted in 5C relating to axon and neuron function or apoptosis and stress, respectively.

**Table S5.**
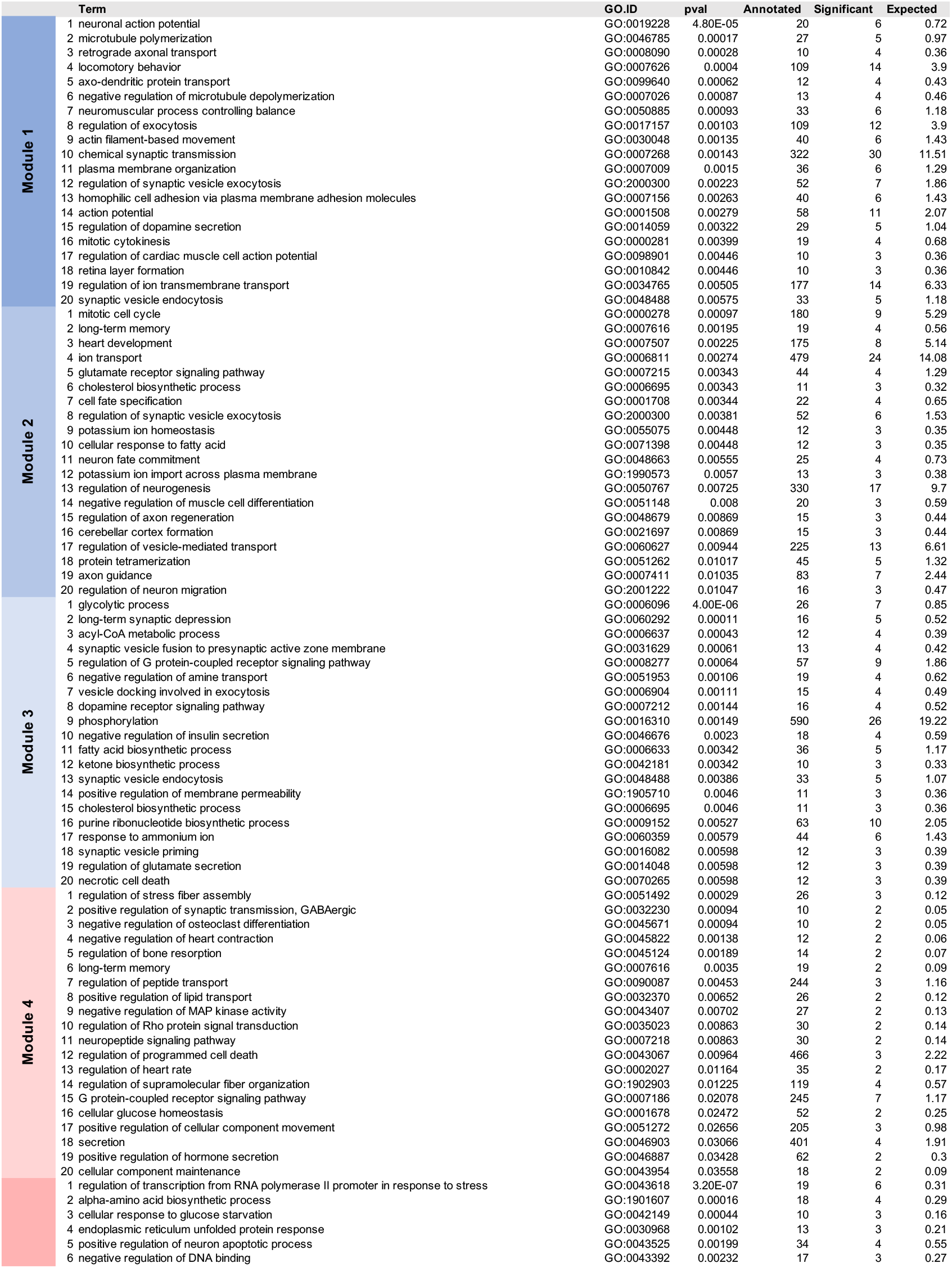
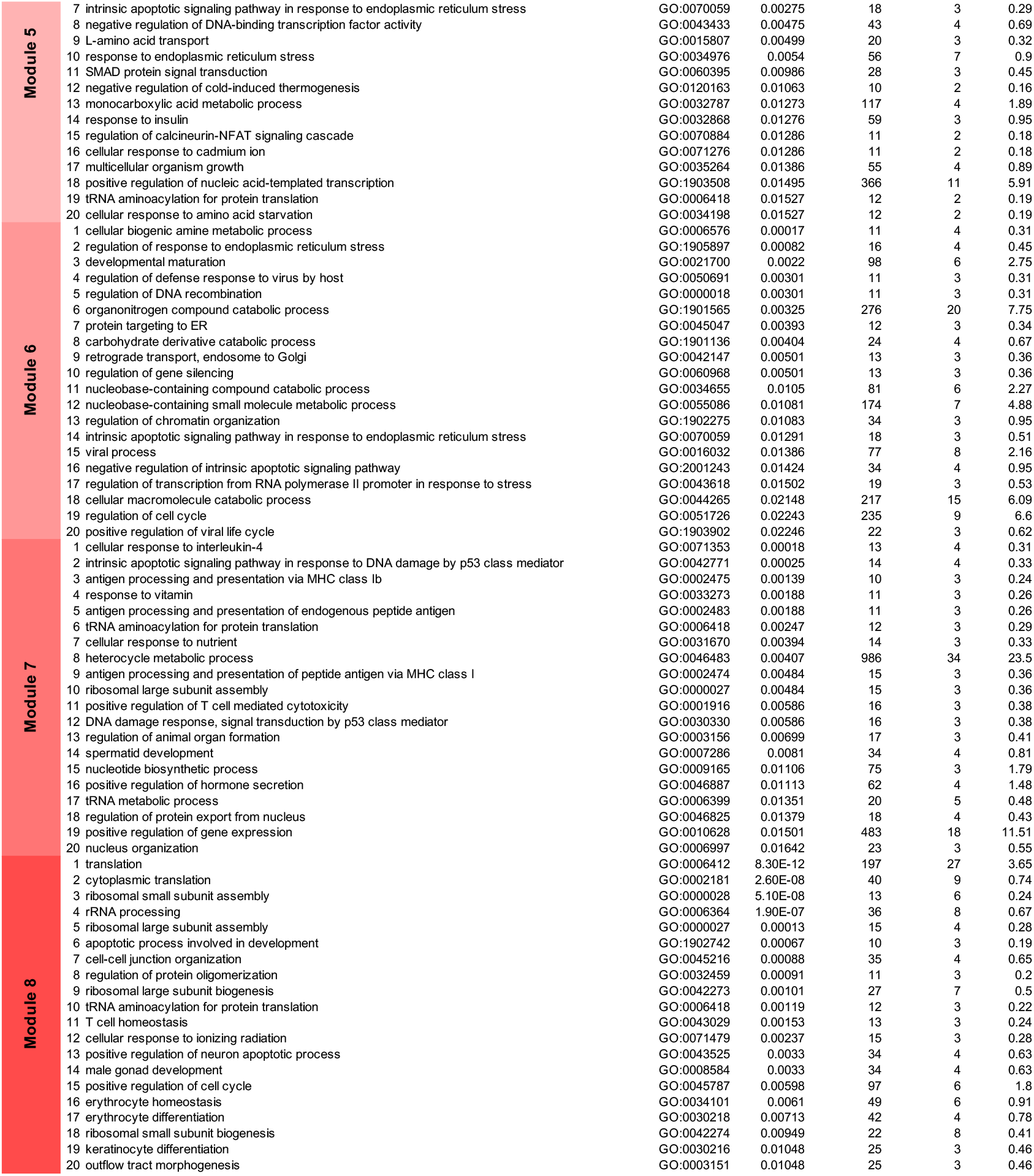
Top 20 significantly enriched biological process GO terms for each gene module (Mod1-8)

**Table S6.1.**
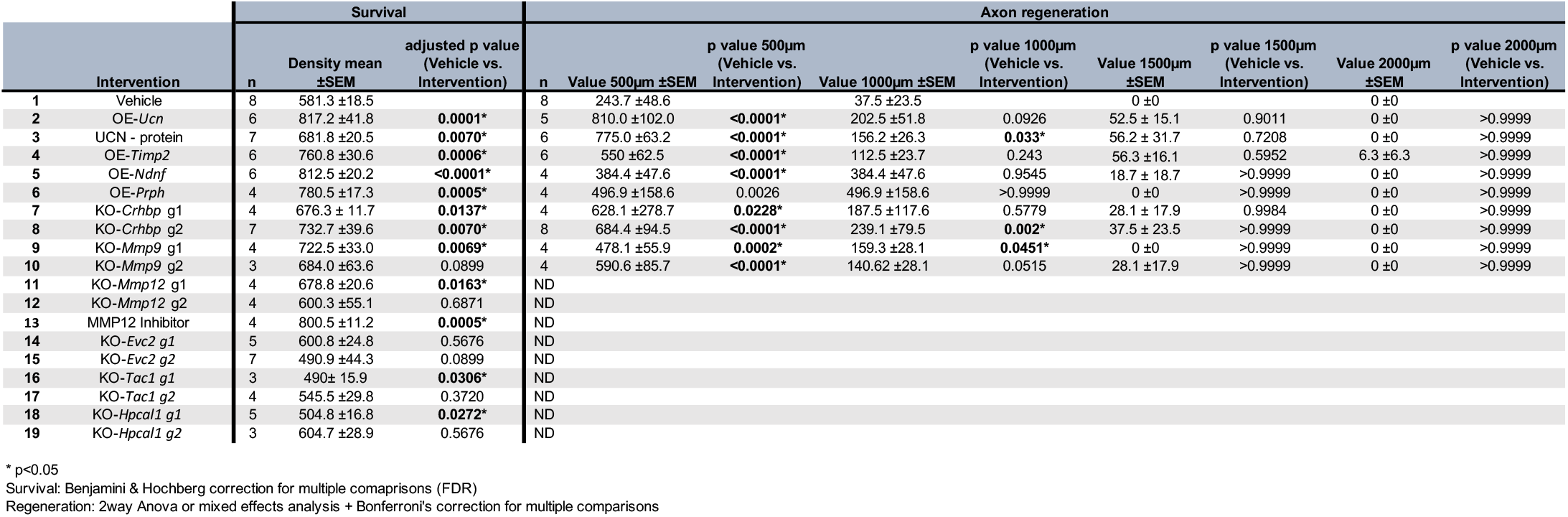
Quantification of RGC survival and axon regeneration at 14dpc

**Table S6.2.** Area under the curve statistics for axon regeneration

| n | Intervention | p value (Vehicle vs.<br>Intervention) |
| --- | --- | --- |
| 1 5 | OE-*Ucn* | **0.0016\*** |
| 2 6 | UCN - protein | **8.1830E-06\*** |
| 3 6 | OE-*Timp2* | **0.0013\*** |
| 4 4 | OE-*Ndnf* | 0.1953 |
| 5 4 | OE-*Prph* | 0.1741 |
| 6 4 | KO-*Crhbp* g1 | 0.0689 |
| 7 8 | KO-*Crhbp* g2 | **0.0061\*** |
| 8 4 | KO-*Mmp9* g1 | **0.0054\*** |
| 9 4 | KO-*Mmp9* g2 | **0.0015\*** |

\* p<0.05
Integral Analysis ("Area under the curve") + followed Student's T-test

**Table S7.**
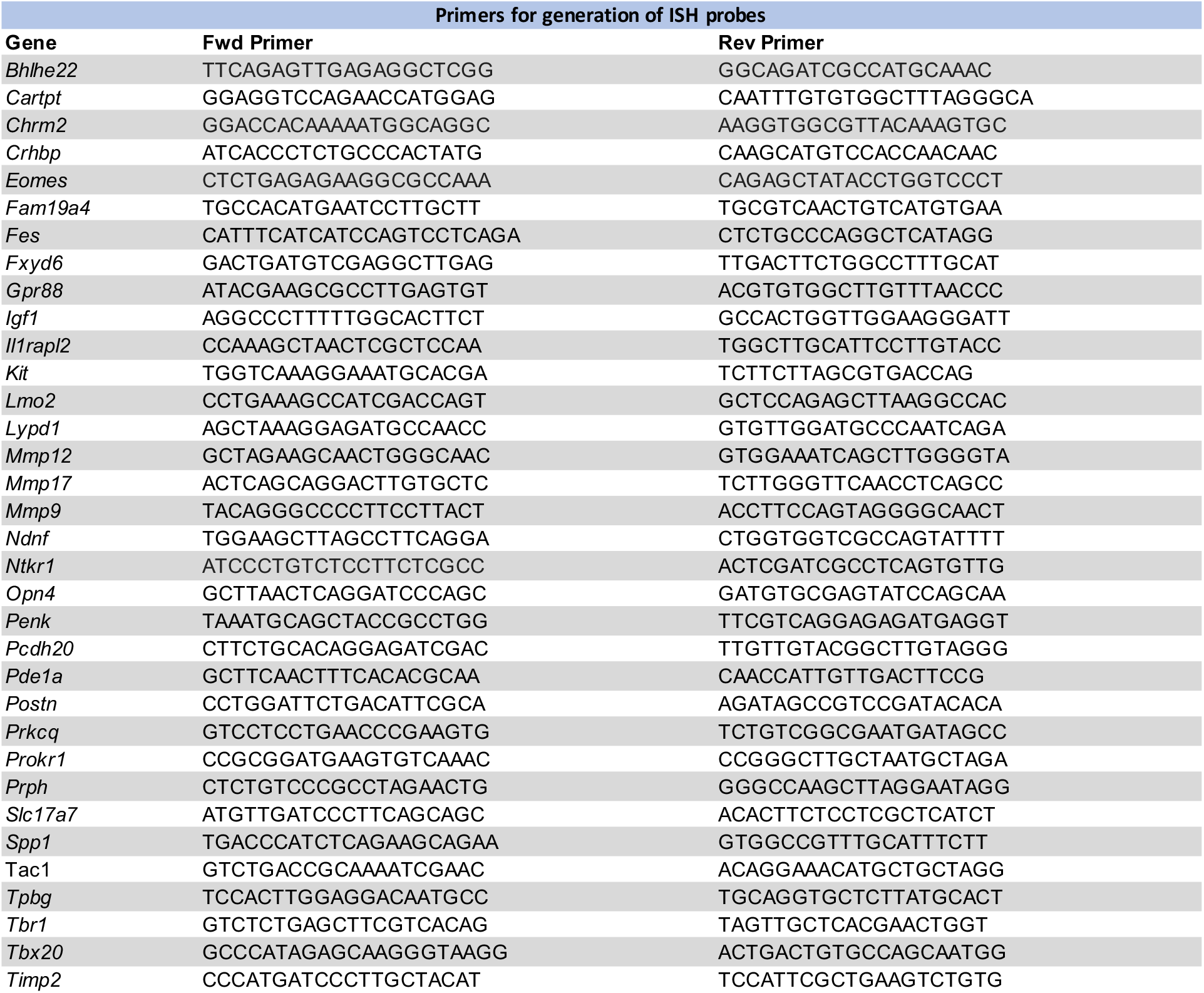
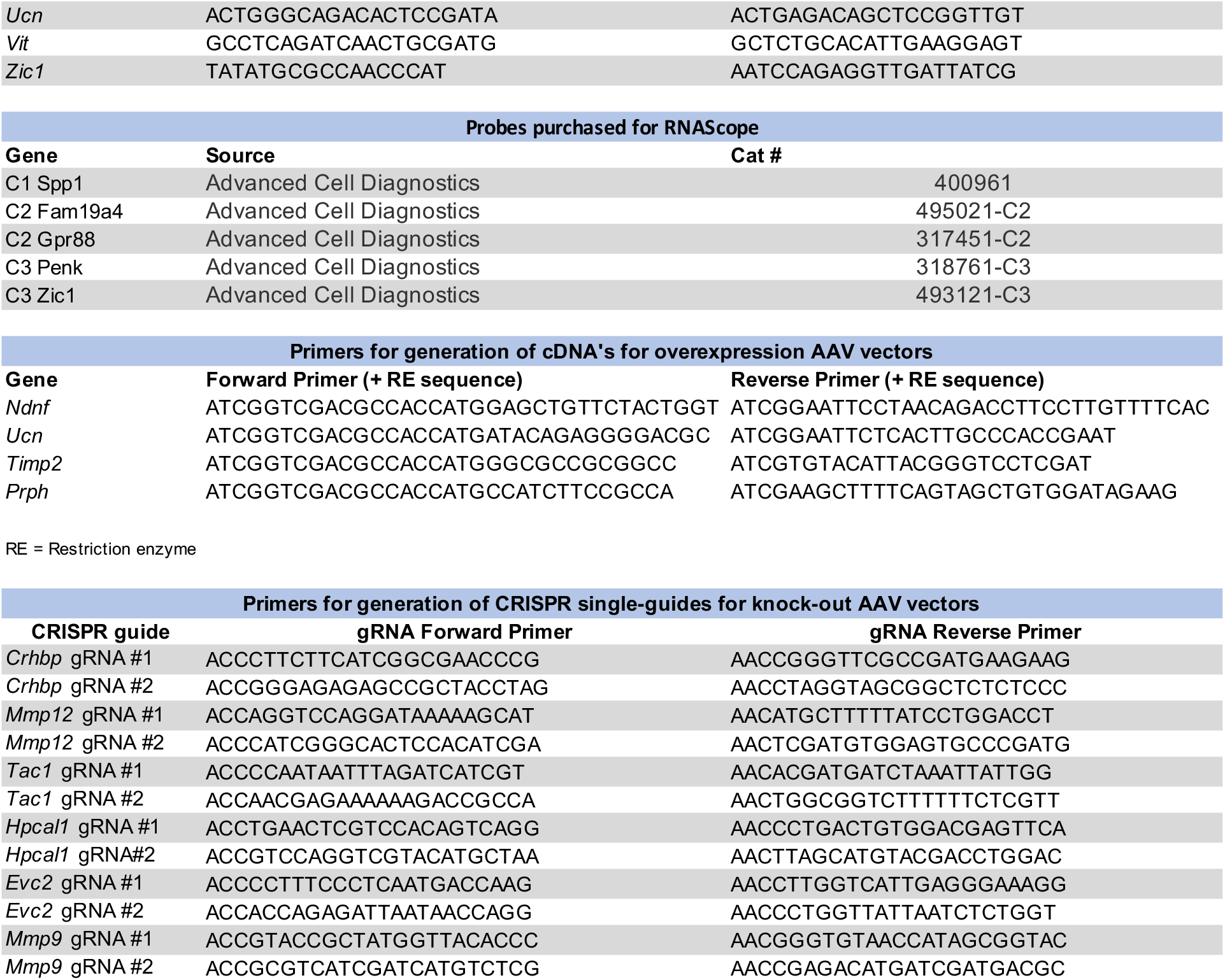
Primers and oligonucleotides used in this paper

## Contact for Resource and Reagent sharing

Further information and requests for resources and reagents should be directed to and will be fulfilled by J.R.S.

## Methods

### Mice

All animal experiments were approved by the Institutional Animal Care and Use Committees (IACUC) at Harvard University and Children’s Hospital, Boston. Mice were maintained in pathogen-free facilities under standard housing conditions with continuous access to food and water. All experiments were carried out in adult mice from 6 to 20 weeks of age. The following mouse stains were used for both fluorescence activated cell sorting (FACS) and histology: *Vglut2-ires-cre (Slc17a6^tm2(cre)Lowl^/J*;(Vong et al., 2011)) crossed to the cre-dependent reporter *Thy1-stop-YFP Line#15 (B6.Cg-Tg(Thy1-EYFP)15Jrs/J* (Buffelli et al., 2003), *C57Bl/6J* (JAX # 000664), *TWY3-YFP* (Kim et al., 2010). The following mouse strains were used only for histology: *Kcng4-cre* (Duan et al., 2015) crossed to *Thy1-stop-YFP Line#1* (Buffelli et al., 2003), *Jam-Creer* (Kim et al., 2008) and *Cck-Cre* (Taniguchi et al., 2011) crossed to *Thy1-stop-YFP Line#15, Pv-Cre* (JAX #017320) and *Opn4-Cre* (Ecker et al., 2010) crossed to Rosa-lox-STOP-lox-Tomato (Madisen et al., 2010), *YFP-H* (Feng et al., 2000), *B6.Cg-Tg(Hlxb9-GFP)1Tmj/J* (AKA Hb9-GFP) (Kim et al., 2010), *TWY7-YFP* (Kim et al., 2010), and *Penk-cre* (JAX #025112). For Crispr-induced gene knockdown experiments *Vglut2-Cre* crossed to the *Rosa26-LSL-Cas9 knockin* (JAX #024857) was used.

### Optic Nerve Crush

After anaesthesia with ketamine/xylazine (ketamine 100-120 mg/kg and xylazine 10 mg/kg), we performed optic nerve injury as previously described (Park et al., 2008). Briefly, the optic nerve was exposed intraorbitally and crushed with fine forceps (Dumont #5 FST) for 2s approximately 0.5-1mm behind the optic disc. Eye ointment was applied post-operatively to protect the cornea.

### Cell preparation and FACS

Retinas were dissected in AMES solution (equilibrated with 95% O2/5% CO2), digested in papain, and dissociated to single cell suspensions using manual trituration in ovomucoid solution. Cells were spun down at 450g for eight minutes, resuspended in AMES+4%BSA to a concentration of 10 million cells per 100μl. 0.5μl of 2μg/μl anti-CD90 (conjugated to various fluorophores) (Thermo Fisher Scientific) per 100μl of cells was incubated for 15 minutes, washed with an excess of media, spun down and resuspended again in AMES+4%BSA at a concentration of ~7 million cells per 1 ml. For Actinomycin-treatment experiments, cell preparation was performed as above and 30μM ActinomycinD (Millipore Sigma) was added to the oxygenated AMES, papain, and ovomucoid solutions. Cells were then resuspended in AMES + 4%BSA + 3μM ActinomycinD. Just prior to FACS the live cell marker calcein blue was added. Cellular debris, doublets, and dead cells (Calcein Blue negative) were excluded, and RGCs were collected based either on high CD90 expression alone, or on CD90 and GFP coexpression. The former was used when tissue came from C57Bl6/J mice, and the latter when *Vglut2:cre;Stp15* mice were used. Cells were collected into ~100ul of AMES+4%BSA per 25,0 sorted cells. Following collection cells were spun down and resuspended in PBS+0.1% non-acetylated BSA at a concentration range of 500-2000 cells/ul for droplet-based scRNAseq per manufacturer’s instructions (10x Chromium). YFP+ RGCs from *TWY3-YFP* were collected in the same way as RGCs from *Vglut2:cre;Stp15* mice, but single cells were sorted into 96 well plates as described below.

### RNA-sequencing

#### 3’ droplet-based scRNA-seq

Single cell libraries were prepared using the Single-cell gene expression 3’ v2 kit on the Chromium platform (10X Genomics, Pleasanton, CA) following the manufacturer’s protocol. Briefly, single cells were partitioned into Gel beads in EMulsion (GEMs) in the Chromium instrument followed by cell lysis and barcoded reverse transcription of RNA, amplification, enzymatic fragmentation, 5’ adaptor attachment and sample indexing. On average, approximately 8,000-12,000 single cells were loaded on each channel and approximately 3,000-7,000 cells were recovered. Libraries were sequenced on NextSeq 500 or Illumina HiSeq 2500 platforms (Paired end reads: Read 1, 26 bases, Read 2, 98 bases).

#### W3-GFP Smart-seq2

We profiled 768 single cells from the TYW3 mouse line (Kim et al., 2010), in which a subset of RGCs with S3-laminating dendrites is labeled with YFP. We sorted single YFP+ cells from dissociated retinas into wells of 96-well plates preloaded with 2 μl of lysis buffer containing 0.5% NP-40, 10 mM Tris-HCl, pH 7.6, 0.1 mM EDTA, 2 U/μl RNase Inhibitor (Clontech/TaKaRa). We generated single cell RNA-Seq libraries using a modified Smart-seq2 method (Ding et al., 2019; Picelli et al., 2013) with the following minor change: We added 3 μl instead of 4 μl of master mix containing only 1.7 μl instead of 2.7 μl of 1 M Trehalose (Sigma-Aldrich) directly to the 2 μl cell lysate without a SPRI bead cleanup step. We pooled and sequenced the libraries with paired-end reads (50 bases for read 1 and 25 bases for read 2) on two flowcells with a NextSeq 500 instrument (Illumina).

### Histological Methods

Eyes were either collected from animals intracardially perfused with 15-50ml of 4% paraformaldehyde (PFA), and post-fixed for an additional 15 minutes, or dissected from an nonperfused animal and immersion fixed in 4% PFA for 30 minutes. Eyes were transferred to PBS until retinas were dissected, following which retinas were either used for wholemount IHC or sunk in 30% sucrose and embedded in tissue freezing media to cryosection into 20-25μm thick crosssections. To immunostain retinal wholemounts, retinas were incubated in protein block (5% normal serum, 0.3% triton-x, 1x PBS) for 3-14 hours, followed by incubation with primary antibodies (in protein block) for 5-7 days in block, and secondary antibodies (in 1x PBS) overnight. All incubations were done at 4°C with gentle rocking. Retinal sections were then used for IHC or fluorescent *in situ* hybridization (FISH). For IHC, slides were incubated for 1 hour in protein block, primary antibody incubation overnight, and secondary antibodies for 2-3 hours. Initial block and secondary antibody incubation were done at room temperature and primary antibody incubation at 4°C. Probe generation and FISH was performed as described previously with minor modifications (Shekhar et al., 2016), specifically a reduced digestion with Proteinase K (0.5ug/ml for 5 minutes) to preserve the integrity of the GCL. In some cases, FISH was performed using the commercially available RNAscope fluorescent multiplex assay according to manufacturer’s instructions, with minor modifications (ACDbio). Specifically, we excluded the step of boiling slides in target retrieval solution, which disrupted IHC staining.

### Intravitreal injections for manipulation experiments

For AAV-based experiments, mice were anaesthetized with ketamine/xylazine (ketamine 100-120 mg/kg and xylazine 10 mg/kg) and injected intravitreally with ~2μl of volume of AAV2 (in 1x PBS) carrying the gene of interest driven by a CAG promoter, or an sgRNA driven by a U6 promoter, two weeks before crush. Concentration of viruses was adjusted to ~5 x 10^12^. Urocortin (rat) protein (Millipore Sigma, ~2μl of 40μM in 1x PBS + 0.1% acetic acid) was injected intravitreally at 2dpc. Mmp12 inhibitor (Mmp408) (Millipore Sigma, ~2μl of 2mM in 1x PBS+1:20 DMSO) was injected intravitreally at 2, 5, 8, 12dpc. For injections, we first removed ~2μl intravitreal fluid from the eye with a sterile micropipette glass. A sterile micropipette glass tip or 33-gauge Hamilton syringe was then inserted through the sclera about 0.5 mm posterior to the limbus and into the vitreal chamber without touching the lens and delivered reagent (~2μl) was injected through the same injection site. After injection, antibiotic ophthalmic ointment was applied and mice stayed warmed on a heating pad until fully awake.

### Design of overexpression and knockdown vectors

Boston Children’s Hospital Viral Core provided AAV virus. The AAV2-based Crispr/Cas9 approach we employ here has been established as an effective modality for somatic knockdown in adult mouse RGCs (Hung et al., 2016). To account for possible off target effects, we tested two gRNAs per gene, and for added RGC-specificity, we delivered AAV2 single-guide RNA (sgRNA) expression vectors to the eyes of mice that express Cas9 specifically in RGCs (VGlut2-Cre; LSL-Cas9-eGFP), which lead to high infection rates as exemplified here for *Crhbp* sgRNA #2 and OE Ucn (**Figure S7A**). Vectors and sequences used for manipulation experiments are displayed in Key Resources Table and **Table S7**.

### Anterograde tracing of regenerating axons

To assess axon regeneration, the axons were anterogradely labeled by injecting CTB conjugated with Alexa-647 (Life Technology) via an intravitreal injection 48 hours before sacrifice. After 4% PFA perfusion mice heads were postfixed for 3 hours in 4% PFA. Ooptic nerves were microdissected and meninges surrounding the nerve were removed. Nerves were then cleared by the protocol provided from Visikol. Nerves were dehydrated with 100% methanol for 4 minutes and then transferred into Visikol Histo-1 solution for overnight incubation at 4°C. The next day the nerves were incubated in Visikol Histo-2 solution for at least 2 hours before mounting them in Visikol Histo-2 solution and imaged with the LSM710 confocal microscope. Optic nerves showing incomplete crushes as evidenced by continuous labeling of axons through the chiasm and/or a different morphology then regenerating axons (pearls on a string) were excluded from the analysis; they comprised <4% percent of nerves analysed.

### *In vivo* Electrophysiology

#### Implantation of mesh electrodes

Fabrication and non-coaxial implantation of the mesh on the retina surface have been reported by (Hong et al., 2018) except that the mesh electronic probe produced for our experiments carried 32 independently addressable recording electrodes rather than 16 electrodes in the previous report. The mesh was loaded into a sterile borosilicate capillary needle Inner diameter: 200μm, outer diameter: 330μm; Produstrial LLC, Fredon, NJ). Mice were anesthetized by intraperitoneal injection of a mixture of ketamine/xylazine (ketamine 100-120 mg/kg and xylazine 10 mg/kg), the optic nerve of the right eye was crushed as described above and immediately after that a sterile 27-gauge hypodermic needle (BD Technologies, Durham, NC) was used to puncture a hole for sclerotomy below the limbus at the lateral canthus for guiding the insertion of the capillary needle and at the medial canthus for draining the injected liquid to reduce intraocular pressure during injection. Using a stereotactic stage, the capillary needle loaded with the mesh was allowed to advance through the pre-punctured hole at the lateral canthus until its tip reached the nasal part of the retina, taking special caution to avoid damaging the lens. Controlled injection of the mesh electronic neural probe to achieve precise placement was previously reported (Fu et al., 2016; Hong et al., 2018; Hong et al., 2015). After a 2-3mm length of the mesh has been injected and placed on the retina surface, conventional coaxial injection was used while the capillary needle was withdrawn simultaneously, leaving an external portion of the mesh outside the eye. The exit point of the mesh probe in the lateral canthus was secured with a small amount of Kwik-Sil adhesive silicone elastomer (Word Precision Instruments, Sarasota, FL). The medial canthus of the eye was sealed with 3M^TM^Vetbond^TM^Tissue Adhesive (Santa Cruz Biotechnology Inc., Dallas, TX). Antibiotic ointment was applied after eye injection.

The external portion of the mesh has all input/output (I/O) pads, which were unfolded onto a 32-channel flexible flat cable (FFC, #0150200339, Molex, Lisle, IL) for individually addressable I/O connection (Fu et al., 2016; Hong et al., 2018; Hong et al., 2015). After I/O connection, the bonding region of mesh on the FFC was carefully covered with dental cement to the skull with METABOND®. A mouse head-plate, with an opening, for head-fixation during retina recording and visual field stimulation was also cemented to the skull using METABOND®. Additional dental cement was carefully applied to cover the silicone previously applied at the lateral canthus of the injected mouse eye for protection of mesh electronics without touching any part of the mouse eye, eyelids or the mesh, resulting in a monolithic piece of dental cement protecting the mesh electronics and the FFC, and a chronically stable interface for long-term retina electrophysiology.

#### *In vivo* recording and stimulation protocol

We obtained recordings from mice every one or two days, starting from Day 1 following ONC and mesh injection. Mice were placed in a Tailveiner restrainer (Braintree Scientific LLC., Braintree, MA) with the head-plate secured to reduce mechanical noise during recording and fix the visual field of the recorded eye during visual stimulation. The FFC was connected to the signal amplifier and digitizer (Intan Technologies, Los Angeles, CA) and RGC activities were recorded while visual stimuli were presented to the mouse on a computer screen (20.5”×12.5”), placed a distance of 20cm from both eyes of the mouse, covering an azimuth angle range of ±52°, similar to previously reported protocols (Hong et al., 2018). Recordings were made from the following visual stimuli:

1. *Full-field ON/OFF stimulation:* A full-field projection of a black screen with 4s duration was followed by a full-field projection of a white screen, with its leading edge entering the screen in eight different directions. The full-field projection of the white screen also lasted 4s, which was followed by another 4s full-field projection of black screen with its leading edge moving in the same direction as the preceding white screen leading edge. Each direction was repeated 10 times.
2. *Moving gratings:* Gratings comprising alternating white and black bars filling the entire computer screen and moving in different directions were programmed in MATLAB. A complete moving grating test comprised 10 repetitive trials, where each trial comprised 8 different directions in a randomized sequence. Baseline was established between repetitions with full-field grey screen, which had the same luminous flux as the alternating white-and-black bars. Gratings moved into each direction for 4 seconds. Data was acquired with a 20-kHz sampling rate and a 60-Hz notch filter.

For recordings during both visual stimulation protocols, data was acquired with a 20-kHz sampling rate and a 60-Hz notch filter.

### Computational Methods

#### Alignment and quantification of gene expression in 3’ droplet-based scRNA-seq data

Demultiplexing and alignment of sequencing reads to the mouse transcriptome (see below) was performed using the Cell Ranger software (version 2.1.0, 10X Genomics). For each sample (i.e. 10X channel/reaction), Cell Ranger generated a matrix of gene counts across cells. We used the option ‘‘--forcecells 6000’’ in “cellranger count” to deliberately extract a larger number of cell barcodes in the data, as we found that the automatic estimate of Cell Ranger was too conservative. Here, 6,000 represented a ‘‘loose’’ upper bound on the number of cells that could be recovered, a value that was calculated from the measured density of the cell suspension loaded onto every channel per the manufacturer’s guidelines.

The 10 samples that were used to assemble the adult RGC atlas (**Figure 1**) were profiled prior to ONC experiments, and were aligned to the standard mm10 mouse transcriptomic reference that is included in the Cell Ranger suite. For ONC-related scRNA-seq experiments, we aligned the sequence data to an updated transcriptomic reference (see below). The count matrices corresponding to the atlas and the ONC datasets were combined separately to generate consolidated matrices *C_ij_* representing the Unique Molecular Identifier (UMI)-based transcript counts for gene i in cell j. For normalization, we first divided each *C_ij_* by 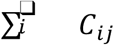 representing the library size of cell j to obtain a “concentration” matrix 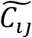. We then multiplied 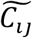 by the median library size M 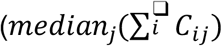 to obtain the transcripts per median (*TPM_ij_* matrix. We defined the normalized expression matrix *E_ij_* = log (*TPM_ij_* + 1).

#### Updated transcriptome for crush experiments

We obtained high quality total RNA from uninjured control retinas and from a pooled sample of 1, 2, and 14dpc retinas, and processed these separately to generate two (Ctrl and ONC) strand-specific RNA-seq libraries using the TruSeq Total RNA kit (Illumina Inc.). Each library was sequenced on a NextSeq 500 system to obtain 75 million 100bp paired end reads. Next, we used the two datasets to assemble a new mouse retina-specific transcriptomic reference, beginning with the original mm10 mouse reference as a scaffold. We followed a procedure similar to the one we used recently to update a macaque retina transcriptome (Peng et al., 2019). Briefly, following published guidelines (Pertea et al., 2016), we initially mapped the TruSeq reads onto the mm10 transcriptome in a strand-specific manner using the Hisat2 software with command line options “--dta--rna-strandedness RF”. Next, we used StringTie v1.3.3 (Pertea et al., 2016) with command line option “--rf” to assemble a new transcriptome based only on the TruSeq reads. After the initial assembly, we reran StringTie with the command line option “--merge” to unify the assembled transcripts with the previous reference to obtain an updated transcriptome annotation. This resulted in modified definitions of transcripts existing in the original reference, as well as novel transcripts supported by the TruSeq reads. While the modified transcripts retained gene names from the original annotation, the novel transcripts were initially named according to Stringtie’s naming convention (e.g., MSTRG.7121).

#### Defining a molecular atlas of mouse RGC types

Our scRNA-seq libraries consisted of three biological replicates (Batch 1-3). In each case, we collected CD90+GFP+ cells from the retinas of Vglut2-cre;Stp15 mice. Cells in Batch 1 were processed in two 10X channels, while cells in Batches 2-3 were processed in four channels each, with an estimated recovery of ~4000-5000 cells per channel. We consolidated the expression matrices, and selected cells where a minimum of 1200 genes were detected. This resulted in a total of 39,750 cells across the three biological replicates (8,091 cells from Batch 1, 17,327 cells from Batch 2 and 14,332 cells from Batch 3).

We identified 1285 highly variable genes {*g_i_*} in the data using our previously described Poisson-Gamma model (Pandey et al., 2018) based on the raw count matrix *C_ij_*, and used the filtered expression matrix *E_{gi}j_* for further analysis. We used randomized PCA (Halko et al., 2011) to reduce the dimensionality of the data, and identified 61 statistically significant PCs, identified based on comparison of the empirical eigenspectrum with a “null” spectrum based on the Tracv-Widom distribution, as described earlier (Peng et al., 2019). Next, we built an unweighted k-nearest neighbor (k-NN) graph (k=40) on the cells based on euclidean distance in the 61dimensional PC space. Each edge connecting a pair of cells *i* and *j* was re-weighted using the Jaccard-metric of neighborhood similarity,

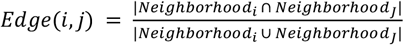

Where *Neighborhoodi* denotes the cells that are nearest neighbors of cell *i*, and |x| denotes the number of elements in the set denoted by *x*. Using the Infomap algorithm (Rosvall and Bergstrom, 2008), which in our previous experience over-clusters the data (Shekhar et al., 2016), we identified 304 clusters in the data with sizes ranging from 502 cells to 6 cells. We scored each cluster based on cell class-specific signatures for the major retinal neuronal classes – RGCs, Amacrine cells, Photoreceptors, Bipolar cells, Muller glia, horizontal cells, pericytes and microglia (**Table S1**). Removal of contaminant clusters (Amacrine Cells, Photoreceptors, Immune Cells and Glia) resulted in 251 RGC clusters comprised of 35,699 cells.

While the sensitivity of Infomap was beneficial in identifying, and eliminating contaminants, a large number of the remaining RGC clusters could not be justified based on differential gene expression. In particular, a hierarchical clustering analysis (not shown) revealed a number of instances where multiple clusters were close together in transcriptional space that exhibited minor and graded differences in gene expression, likely representing oversplitting of a single RGC type/state or batch effects. We therefore reclustered the RGCs using the Louvain algorithm (Blondel et al., 2008) applied on a Jaccard-weighted kNN graph (k=30), and applied the iterative approach described earlier (Peng et al., 2019) to identify 45 molecularly distinct clusters (putative RGC types). Through manual inspection of a few Louvain clusters, we confirmed that they were supersets comprised of Infomap clusters that were transcriptionally proximal.

We bootstrapped on the number of genes and number of PCs to ensure that our results were insensitive to variations in these choices (not shown). To further assess the robustness of these clusters, we used the output of two alternatives to PCA to reduce the dimensionality of the data prior to Louvain Clustering – Independent Component Analysis (ICA; (Comon, 1994)) and Liger (Welch et al., 2019), a recently proposed non-negative matrix factorization (NMF; (Lee and Seung, 1999)) based technique based data integration method. Encouragingly, the clusters identified in either space were highly consistent with the results from PCA (Figure S1C,D).

#### Identifying 2-marker combinations to label RGC types in the atlas

We first identified genes enriched in each RGC type relative to the rest using the MAST framework (Finak et al., 2015). In contrast to our previous study on retinal bipolar cells (Shekhar et al., 2016), we found that most RGC types were not specifically labeled by single genes. We therefore evaluated combinations of two genes on their ability to specifically label cells of a given type.

For each type, we first selected genes that individually showed either a >2-fold enrichment (“+ve markers”) or < 2-fold depletion (“-ve markers”) compared to the background of other types. Here, fold-enrichment or depletion of gene *i* was evaluated as the max(*r*_1_, *r*_2_) where,

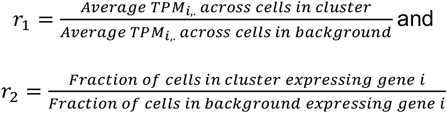

Using these genes, for each RGC type we evaluated (1) all pairwise combinations of +ve markers, and (2) all pairwise combinations consisting of one positive and one negative marker for their ability to specifically label the type based on the area under the precision recall curve (AUCPR). AUCPR is a measure of cluster specificity as described earlier (Pandey et al., 2018). We specifically favored marker combinations that had a higher precision (low proportion of false positives) compared to recall (low proportion of false negatives), as false negatives could arise because of dropouts in scRNA-seq data. The top candidates are displayed in **Figure 1F**. For few types, we found that it was necessary to screen three marker combinations for achieving higher precision.

#### Assigning injured RGCs profiled following ONC to types using iGraphBoost

The ONC dataset consisted of cells enriched for RGCs profiled at 6 time points following crush (0.5, 1, 2, 4, 7, and 14dpc), and a separate control dataset (0dpc) independent of the Atlas dataset. We first used an Infomap-based clustering procedure (see above) to remove non-RGCs from the data. he iGraphBoost procedure used to assign injured RGCs to types following ONC was as follows:

Beginning with the RGC atlas as training data, iGraphBoost proceeds by sequentially assigning cells to types at each time point through supervised classification followed by graph-based voting (see below), beginning with 0dpc. Cell type assignments at each timepoint in the ONC dataset are incorporated into a “time point-specific” atlas, which, combined with previous atlases, is used to classify cells at the next time point. This allows us to disambiguate changes in cell state due to injury from the intrinsic molecular distinctions of each RGC type.

Given RGCs atlases at times *t*_0_, *t*_1_,...,*t_i_*, we describe here the procedure to assign RGCs at time *t*_*i*+1_ to types. Here, *t*_0_ may be regarded as the control RGC atlas and *t*_1_ may be regarded as 0dpc. We train decision-tree based ensemble classifiers Ω_t_ at each time *t* < *t*_*i*+1_. We tested Random Forests (R package *randomForest*) and Gradient-boosted trees (R package *xgboost)*, both yielding consistent results. Results presented were derived from Gradient-boosted trees. Each classifier Ωt was trained on 70% of cells at time *t*, and validated on the remaining 30% “held out” cells. The classifier labels were compared to the true labels of the “held out” cells to compute precision and recall values for each of the 45 RGC types. In the assignment of cells at *t*_*i*+1_, a classifier Ω_t_’s vote for a class *m* (when *p^m^_k,t_* ≥ 0.8 was only “trusted” if during the validation round both the precision and recall for class *m* exceeded 0.9.

In **Step 1**, we begin by individually applying the classifiers Ω_t_ to each cell *k* at time *t*_*i*+1_. When applied to cell *k* Ω_t_ assigns a probability vector 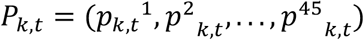 where *p^m^_k,t_* is the Ω_t_ assigned probability that cell *k* belongs to RGC type *m*. *p^m^_k, t_* is the fraction of decision trees in Ω_t_ that vote for class *m*. For the classifier Ω_t_, we assign cell *k* to class *m^t^* = *argmax_m_p^m^_k,t_* if and only if 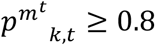, and Ω_t_ can be trusted with regards to class *m^t^*(see above). If 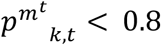, we regard cell *k* as “unassigned” from the point of view of Ω_t_. The final assignment of cell *k* in Step 1 is determined by collectively considering the votes from all the Ω_t_ such that cell *k* is assigned to type *m* if and only if at least one classifier among Ω_t_ assigns cell *k* to RGC type *m*, and the remaining classifiers deem it “unassigned”. In cases where two or more classifiers among Ω_t_ assign cell *k* to different RGC types, it is deemed “unassigned”.

Step 1 resulted in a subset of RGCs at time *t*_*i*+1_ being assigned to types with high confidence. In **Step 2**, we use these as “anchors” to propagate labels using the neighborhood relationships. We built a k-nearest neighbor graph on the RGCs with k=15 at *t*_*i*+1_ in PCA space, which connects cells based on their transcriptional proximity. We hypothesized that if an unassigned cell’s neighbors were predominantly of a single type, this can be used as evidence to assign the transcriptional identity of the unassigned cell. Thus, we iteratively loop through the unassigned cells from Step 1 and assign a cell *k* to type *m*, if > 67% of the *k*=15 nearest neighbors of cell *k* are of type *m*. Each iteration decreases the fraction of unassigned cells, and the procedure terminates if the fraction of unassigned cells decreases less than 0.5%. The results were insensitive in variations of the voting threshold between 50% and 80%, although there was a drop in quality outside of these limits. A high voting thresholding (~90%) resulted in a number of unassigned cells, whereas a low voting threshold resulted in a number of spurious assignments, as assessed by post hoc DE analysis. We typically conducted 2-3 “passes” of nearest-neighbor voting, successively propagating labels in the data, stopping when the proportion of newly classified cells was dropped below 1%.

Thus steps 1 and 2 assign type identities RGCs at *t*_*i*+1_. While unassigned cells remained, they were far fewer than the naive “one-step” classification approach, involving a direct assignment of cells based on a classifier Ω_t0_ trained on the atlas (**Figure S3**).

#### Clustering and visualization of atlas and injured RGC transcriptomes

As described previously, we used the PCA coordinates of atlas RGCs to determine clusters using the Louvain-Jaccard algorithm. RGCs were visualized on a 2D map using t-distributed stochastic neighbor embedding (tSNE)(Maaten and Hinton, 2008). We used a recently published scalable implementation of tSNE based on fast interpolation (FI-tSNE) (Linderman et al., 2019).

We initially used a similar procedure to visualize injured RGCs, but found that cells co-clustered by time (**Figure S3A**), resulting from injury related changes, and to a lesser extent, batch effects (distinct batches from the same time point co-clustered, suggesting that batch-effects were not predominant). We therefore aligned injured RGCs using Liger (Welch et al., 2019), a recently proposed non-negative matrix factorization-based algorithm, to disambiguate shared (RGC type-specific signatures) and dataset-specific (injury related changes) features across the different time points. We used the reduced dimensional coordinates provided by Liger as input for FI-tSNE as well as Louvain-Jaccard clustering (**Figure S3A**). Compared to tSNE and clusters computed on PC scores, Liger coordinates and clusters were driven far less by time (days post crush or dpc; **Figure S3B**) compared to iGraphBoost-assigned cell type identity, suggesting that Liger successfully aligns cell types across time. Note that iGraphBoost results did not inform the Liger visualization and clustering).

#### Comparing relative frequencies of RGC subsets between scRNA-seq and immunohistochemistry

We quantified the frequency of multiple combinatorially labeled subsets of RGCs in retinal whole mounts using immunohistochemistry, and compared them to analogous frequencies quantified using the scRNA-seq data at 0 and 14dpc datasets (Table S3). Frequencies were compared to IHC results, which were quantified as described below.

To compute corresponding frequencies in scRNA-seq, we identified RGC clusters that expressed the genes represented in the IHC combination, and computed their relative frequency in the data at the same time point. For example, to calculate the frequency of *SPP1^+^OPN4-cre^-^* labeled cells in scRNA-seq at 14dpc, we identified all RGC clusters that expressed *Spp1*, but did not express *Opn4* at 14dpc. A cluster was regarded as positively expressing a particular gene if >30% of its cells expressed that gene. For IHC combinations involving a transgene, we used either prior knowledge (e.g. W7) or the analysis reported in this paper (e.g. W3) to identify labeled types in the scRNA-seq. We included the “unassigned” RGCs in the background while computing relative frequencies at any time point. **Table S3** describes for each IHC combination, the list of associated types and their corresponding relative frequencies in IHC and scRNA-seq. The procedure to convert estimates of relative frequencies to fractional survival is described below.

#### Quantifying the survival and kinetics of loss of each type following ONC

Following iGraphBoost, RGCs in the ONC dataset was either assigned to one of 45 types, or were labeled “unassigned”. *RF_m_*(*t*), the relative frequency of RGC type *m* amongst surviving RGCs at time *t* was defined as *RF_m_* Here, *N_m_*(*t*) was the number of RGCs assigned to type *m* at time *t*, and *U*(*t*) was the number of unassigned RGCs at time *t. RF_m_*(*t*) values were used to calculate the relative frequency ratio for each type *m* in **Figure 3F** as 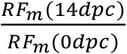

To compute the kinetics of loss of each RGC type, we multiplied the relative frequency *RF_m_*(*t*) of each type with *s*(*t*), defined as the *total* fraction of surviving RGCs at each time *t*, and was estimated independently using IHC (Figure 3I). This enabled us to compute the relative survival (*RS_m_*(*t*)) of each RGC type *m*,

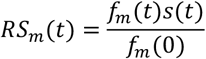

Individual curves in **Figures 3J-L** correspond to *FS_m_*(*t*) for all types. For each type, we obtain a vector 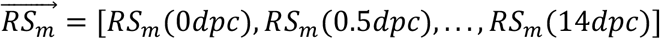, where *RS_m_*(0*dpc*) = 1, by construction. We applied k-means clustering to the vectors 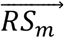, and determined 3 clusters by the elbow method. These three clusters corresponded to the survival groups (**Figure 3M**).

#### Alignment and quantification of gene expression in full length, plate-based scRNA-seq data

For plate-based libraries, expression levels of gene loci were quantified using RNA-seq by Expectation Maximization (RSEM) (Li and Dewey, 2011). Raw reads were mapped to a mouse transcriptome index (mm10 UCSC build) using Bowtie 2 (Langmead and Salzberg, 2012), as required by RSEM in its default mode. On average, 88% (range 75%-92%) of the reads mapped to the genome in every sample. and 55% (range 20%-66%) of the reads mapped to the transcriptome. RSEM yielded an expression matrix (genes x samples) of inferred gene counts, which was converted to TPM (here, defined as transcripts per million) values and then log-transformed after the addition of 1 to avoid zeros. After filtering cells with low QC metrics (< 400,000 mapped reads, transcriptomic mapping rate < 35% and < 1500 genes detected), we selected 636 cells for further analysis.

#### Analysis of W3 RGCs

An initial clustering analysis using Louvain-Jaccard clustering on a kNN graph (k=15) in PCA space (17 significant PCs) identified 7 groups of cells comprised of RGCs (n=341), microglia (n=51), bipolar cells (n=9), amacrine (n=67), rods (n=73), doublets involving RGCs and rods (n= 77), and low quality cells (n=18). Next, we separately reclusterd the 341 RGCs using a similar procedure (13 significant PCs), and found 6 clusters. Supervised classification analysis using random forests trained on the atlas data matched 5/6 clusters 1:1 to atlas clusters C2 (W3D1), C4 (F-mini-OFF), C3 (F-mini-ON), C6 (W3B), and C23 (W3D2). The 6 SS2 cluster mapped to C30 (W3D3) and C21 (Tbr1-S2), respectively.

#### Definition of resRGCs, susRGCs and intRGCs

resRGCs were initially defined as those RGC types *m* for which *RFR_m_* > 2, which included 7 types (C10, C22, C31, C33, C40, C42, C43; see **Figure 3E**). Independently, as described above, we applied k-means clustering on the vectors 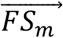, which resulted in the grouping of the 45 RGC types into 3 survival groups. The first cluster comprised the 7 resRGC types. A second cluster comprised 27/45 vulnerable types whose survival rapidly declined by 4dpc, and we called these “susceptible RGCs” or susRGCs. The other group, consisting of 11/45 types, also exhibited poor survival at 14dpc, but declined more slowly, and were termed “intermediate RGCs” or intRGCs.

#### Identifying shared differentially expressed genes following ONC, and gene modules

To identify genes that changed significantly across all RGC types, we computed for each gene *g* its average expression 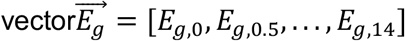, where *E_g,t_* represents the average expression strength of gene *g* across all RGCs at *t* dpc. We defined *E_g,t_* = *P_g,t_ avgTPM_g,t_*, where *P_g,t_* represents the fraction of RGCs at time *t* that express the *g*, and *avgTPM_g,t_* is the average normalized expression of gene *g* in *t* dpc RGCs. We excluded genes such that *min*(*P_g, t_*) < 0.2 and 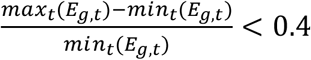, resulting in a total of 3,231 genes.

Next, we randomized the data by shuffling the temporal identities of all RGCs (while maintaining proportions), and computed a randomized average expression vector, 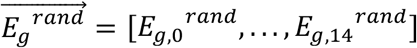. We reasoned that genes with significant temporal variation would exhibit larger differences between 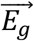 and 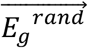, and therefore computed for each gene *g* a deviation score between the actual and randomized expression vectors,

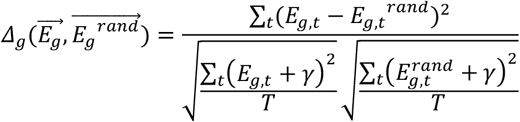

Here, *T*=7, the total number of time points, and *γ*represents a pseudocount which we set to 0.2. Here, the denominator acts as a normalizing factor; in its absence we found that genes with higher expression levels were favored. To evaluate the significance of *Δ_g_* values, we used additional randomizations where we computed 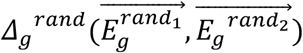 for 10,000 paired randomizations of the data *rand*_1_ and *rand*_2_, which yielded empirical randomized distributions 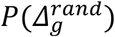 for each gene g. We found that randomized *Δ_g_* values did not exceed 0.2 for any gene. Based on this, we selected genes *g* such that *Δ_g_*> 0.5, which yielded 771 genes. We partitioned these genes into 8 modules using k-means clustering on the 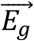 values (elbow method). To identify biological processes enriched in each module, we applied Gene Ontology (GO) analysis using the R package topGO.

#### Identifying genes associated with resilient types as targets for manipulation

To identify genes that were selectively enriched in resRGC types at baseline as well as along the time course, we subsampled the data to equalize the representation of each cluster. This was done to reduce the influence of high-frequency types.

We first compared resRGCs vs intRGCs and susRGCs at baseline using the MAST framework (>2-fold, FDR < 0.0001) to identify genes that were selectively enriched in resRGC types at baseline (**Figure 5F**).

#### P5 to adult correspondence

To evaluate the molecular correspondence between previously published P5 RGC types (Rheaume et al., 2018) and our adult RGC types, we trained graph-boosted trees on our dataset using a set of common variable genes, and assigned each P5 RGC an adult identity. The P5 cluster labels were not used to train the classifier or inform its assignment. We only considered assignments with voting majoring margin > 50% as valid assignments. The correspondence between P5 cluster labels (as in (Rheaume et al., 2018)) and adult type assignments was high, as evaluated by two measures of cluster similarity, the Adjusted Rand Index and Normalized Mutual Information (**Figure S1F**).

### Data analysis of *in vivo* recordings

The electrophysiological recording data was analyzed offline using methods described in previous reports (Fu et al., 2016; Hong et al., 2015). In brief, raw recording data was filtered using non-causal Butterworth bandpass filters (‘filtfilt’ function in MATLAB) in the 250-6000 Hz frequency range to extract single-unit spikes of RGCs. Single-unit spike sorting was performed by amplitude thresholding of the filtered traces, where the threshold was automatically determined the threshold based on the median of the background noise according to the improved noise estimation method (Quiroga et al., 2004). Sorted spikes were clustered to determine the number of RGCs and to assign spikes to each RGCbased on principal component analysis (PCA) using the WaveClus software that employs unsupervised superparamagnetic clustering of single-unit spikes.

For each sorted and clustered spike, a firing time was assigned and all spike firing times belonging to the same cluster (i.e., the same RGC) were used to compute the firing rates in response to different visual stimuli. Analysis of single-unit firing events was different for the two different visual stimulation protocols:

1. Full-field ON/OFF stimulation: Firing rate was computed by dividing the number of firing events of the same RGC over time segments for both ON and OFF phases, averaged over 10 trials. Analysis of variance (ANOVA) was performed using the built-in function ‘anova1’ in Matlab to evaluate the statistical significance between firing rates during ON and OFF phases to determine the light response of each recorded RGC.
2. Moving gratings: The number of firing events for a specific moving direction of the grating was averaged over 10 trials, divided by the duration time for each direction to obtain the average firing rate for all eight directions. Angular distribution of firing rates was plotted in the polar plots to reveal direction and orientation preference and selectivity of the recorded RGCs.

The direction selectivity index (DSi) is defined as:

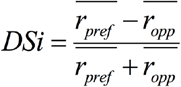

where 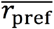 and 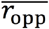 are defined as the average firing rates during moving grating of the preferred direction and that of the opposite direction, respectively.

Similarly, the orientation selectivity index (OSi) is defined as:

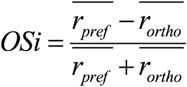

where 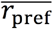 and 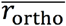 are defined as the average firing rates during moving grating of the preferred orientation and that of the orthogonal orientation to the preferred orientation, respectively.

A DSi or OSi of greater than 0.3 is used to assign a specific single unit to DSGC or OSGC, and that of smaller than 0.3 to non-selective ganglion cells (NSGC).

Cross-correlation analysis of spikes trains assigned to neurons belonging to different channels but showing similar direction/orientation selectivity was carried out to identify potential overlap of recording across channels, that is, the same RGC recorded by more than one electrodes, which was removed from the total count of RGCs.

To determine preferential survival of certain functional types through 14dpc in **Figure 4F-I**, a onesided Fisher’s Exact test was used in R. A p-value smaller than 0.05 were considered statistically significant.

### Quantification and analysis

#### Retinal wholemounts

Immunostained retinal whole mounts were imaged using a Zeiss 710 confocal microscope. For RGC subset quantifications a tile scan of the entire ventral quadrant of the retina was taken to control for topological differences in RGC types. The image was processed in ImageJ (Schindelin et al., 2012) by generating a Z-stack maximum projection of the GCL and by applying a local contrast normalization filter (3 stdev). Circular regions-of-interest (ROIs) (8.3μm in diameter) were placed on >500 RGC somas within a longitudinal area spanning the central-peripheral axis. The centroid position and fluorescent intensity of each ROI was measured in ImageJ. Fluorescent intensity for each marker was plotted against RBPMS (total RGCs) in Matlab using customized scripts and thresholded (3x stdev above linear average of negative cells). Thresholded ROIs were then overlaid onto the wholemount images and visually inspected for accuracy and the density for each possible marker combination was measured. To obtain frequency measurements, the density of each marker was divided by the total density (RBPMS). Individual values are shown in **Table S3**.

For AAV-mediated intervention experiments the maximally infected retinal quadrant was imaged excluding the temporal retina, due to the higher presence of alpha RGCs in this region (Bleckert et al., 2014). Images were processed in ImageJ as previous. Since RGCs at 14dpc are well spaced, it was possible to count them using automated cell segmentation. RGC ROIs were detected using CellProfiler (Carpenter et al., 2006) for the entire retinal quadrant. ROIs were then exported to ImageJ, visually inspected for accuracy, and the intensity and centroid position was measured. To measure marker density, image+ROI overlays were plotted in Matlab and a polygonal boundary region was drawn around the area of the retina that was accurately segmented, taking care to exclude damaged areas, edges where the retina was not viewed en face, and areas with high background staining e.g. blood vessels. Individual values are shown in **Table S6**. To verify the accuracy of this automated segmentation approach, we re-quantified images previously counted by manual ROI placement and found both approaches gave comparable densities.

#### Retinal ISH sections

After ISH, sagittal retinal cross sections through or proximal to the optic nerve (maximal width of retina) were imaged using a Zeiss 710 confocal microscope. To quantify markers, a tile scan image spanning the section was generated for ≥4 sections from ≥2 mice. Marker costaining was counted using the ImageJ cell counter plugin. The fraction of each staining combination was recorded and significance determined by two-tailed student’s t-test (p<0.05).

#### AAV2 transfection rate

Maximum projections of images acquired also for RGC survival quantifications (see above) from AAV2 OE-Ucn (n=2) and AAV2 Crhbp sgRNA #2 (n=7) were quantified via the ImageJ using the “Analyze particles” function (after applying a threshold for background reduction). After visual inspection to ensure accuracy of cell segmentation, the total number of cells labeled with RBPMS (as panRGC marker) and anti-UCN (IHC staining) or Rbpms and mCherry (AAV2 sgRNA for Crhbp tagged with mCherry to allow direct visualization of virus transfection) was assessed to achieve the percentage of RGCs co-expressing either of them.

#### Axon regeneration

The cleared, whole nerve was imaged with a 20X air objective. From the center of the nerve, 7 single stacks (2μm stack size) were maximum projected to a total volume of 14μm per nerve. After defining the crush site, lines spaced equidistant from each other at 500-μm intervals from the crush site to where the longest axon could be detected were introduced for bin-by-bin axon quantification. As described previously (Duan et al., 2015; Park et al., 2008), we quantified the total number of regenerating axons, Σad, by using the following formula: Σad = π*r*^2^ *x* [average axons/mm]/t, where the total number of axons extending distance *d* in a nerve having a radius of *r* was estimated by summing over all sections with thickness *t* (in our case, 14 μm). To assess an overall positive effect on regeneration, we first used a definite integral analysis (= area underneath the curve) and compared each of our interventions to a vehicle control injected samples. Significance is determined by two-tailed Student’s t-test showing p < 0.05 = *. Additionally, bin-by-bin axon quantification was used to assess significant differences between individual distances (500, 1,000, 1,500, 2,000μm) was determined by two-way ANOVA or mixed effects analysis followed by Bonferroni’s multiple comparison test with GraphPad Prism, p < 0.05 = *. Individual values are shown in **Table S6**.

#### Morphometric Analysis

Retinal whole mounts were imaged on an Olympus Fluoview 1000 scanning laser confocal microscope with 20X or 40X oil immersion objectives, optical stacks generated with images taken ever 0.5um. Maximum projections and rotations of images were generated in ImageJ, while brightness and contrast were adjusted in Adobe Photoshop CC. Individual cells were reconstructed and analyzed using the ImageJ plugin Simple Neurite Tracer (SNT, (Longair et al., 2011)). Dendritic size was measured from the area of convex polygons. Dendritic complexity was assessed using total branch points derived from the Stralher analysis in SNT. Significant difference between groups was determined by one-way ANOVA followed by Tukey HSD with SPSS, p < 0.05 = *.

## Data availability

Submission of all the raw and processed datasets reported in this study has been initiated to the Gene Expression Omnibus (GEO) with accession number in process.

